# Small molecule bioactivity benchmarks are often well-predicted by counting cells

**DOI:** 10.1101/2025.04.27.650853

**Authors:** Srijit Seal, William Dee, Adit Shah, Andrew Zhang, Katherine Titterton, Ángel Alexander Cabrera, Daniil Boiko, Alex Beatson, Jordi Carreras Puigvert, Shantanu Singh, Ola Spjuth, Andreas Bender, Anne E. Carpenter

## Abstract

Phenotypic profiling methods, such as Cell Painting and gene expression, have been widely used to predict compound bioactivity, often showing improvement over predictive models based on chemical structures alone. We discovered that a large subset of assays in widely-used benchmark datasets either *directly* relate to cell health and cytotoxicity or are assays intending to capture a more specific phenotype but whose active compounds impact cell count, while inactives do not. As a result, counting cells can achieve similar predictive performance as Cell Painting or gene expression data. Filtering benchmarks to include only assays relating to protein targets reveals that Cell Painting can capture information that cannot be predicted by mere cell counting. We re-evaluated three benchmark datasets used with Cell Painting data and observed that, in many cases, cell count models produced an AUC comparable to models using the full Cell Painting profiles. However, in protein-target-specific benchmarks across 17 distinct protein targets, Cell Painting features demonstrated unique predictive power, outperforming mean balanced accuracy from cell count models with a relative improvement of 19.6%. We propose five practical recommendations for benchmarking machine learning models for predicting bioactivity, including using cell count as a baseline feature. Although multi-class classification applications (such as matching samples based on their morphological profile) are less likely to be predictable by cell count than bioactivity benchmarks, these recommendations are broadly applicable to machine learning for drug discovery.

## INTRODUCTION

Machine learning is now extensively used to predict chemical compounds’ activity in cell-based assays, classifying them as active, inactive, or inconclusive (or toxic, non-toxic, inconclusive).^1^ Such models can reduce the need for expensive laboratory experiments and allow predicting compounds’ efficacy and various types of toxicity early and inexpensively when developing drugs and other chemicals. Predictive models often rely on chemical structures and features derived from them, such as molecular fingerprints, which are easy to generate.^2^ However, improving model performance for compounds outside the training data’s chemical space remains a critical challenge.^3^ Capturing nuances such as activity cliffs—where similar structures show significantly different bioactivity—has proven difficult even for deep-learning architectures, suggesting that chemical representations alone are insufficient for predicting many interactions in complex biological systems.^4^

Phenotypic profiling techniques such as gene expression and morphological profiling have emerged as promising additional data sources for each compound that might augment molecular structures in improving chemical activity prediction. Cell Painting is an image-based profiling method that is high-throughput and cost-effective and, therefore has significant potential.^5^ Cell Painting datasets provide rich phenotypic profiles by applying a range of fluorescent stains that capture the cell’s biological response to various compounds across various cellular compartments and substructures.^6^ This assay has been reported to yield improved performance in downstream tasks compared to chemical structures alone.^5,6^ Since the first demonstration that an image-based assay could predict seemingly unrelated assay outputs^7^, a trend has emerged: using biological response descriptors, such as transcriptomics data or Cell Painting readouts, independently or combined with chemical fingerprints, to enhance predictive models.^5^

For example, Hofmarcher et al. showed that convolutional neural networks trained on Cell Painting features predicted the outputs of 32% of 209 biological assays with area under the curve (AUC) values exceeding 0.9.^8^ Moshkov et al. evaluated chemical structures, Cell Painting images, and gene expression profiles (from the L1000 assay) across 270 assays, finding that each modality independently contributed valuable information.^9^ Fusing the three data modalities improved AUC for 62% of assays (167 out of 270), compared with using chemical structures alone. Sanchez-Fernandez et al. introduced CLOOME, a contrastive learning framework enabling cross-modal querying of Cell Painting images with chemical structures.^10^ They also assessed the transferability of learned representations using linear probing on 209 assay activity prediction tasks with pre-trained image embeddings from CLOOME, which did not use activity data during training. The model achieved an average AUC of 0.714, outperforming fully supervised methods. Ha et al. benchmarked learning models using Cell Painting profiles to predict outcomes of 201 assays, an updated version of the Hofmarcher dataset.^11^ When predicting for assays not included in training the models, they found models performed well across the novel prediction tasks, achieving AUC values ranging from 0.70 to 0.87 as the dataset size increased, effectively generalizing to new assays.

However, when reviewing the assays most accurately predicted in these prior studies, we observed that many were linked to cytotoxicity or cell proliferation. There is a complex interplay between the endpoint one aims to imeasure (or model) in an assay and various confounding factors, which might lead to misleading performance of any predictive models trained on the data. While cytotoxicity/viability are sometimes the desired assay endpoint, these assays are often run side-by-side with an actual assay of interest (in the same cell type for example) to improve assay accuracy by identifying and excluding compounds exhibiting cytotoxic effects. Combining results from the primary assay and the cytotoxicity viability screen enables identifying compounds with specific biological effects, minimizing false positives and off-target effects. For example, Esher et al. addressed the challenge of distinguishing specific from nonspecific (cytotoxicity-triggered) reporter gene activation in six Tox21 high-throughput screening assays.^12^ “Cytotoxicity burst” refers to the activation of stress responses at compound concentrations near the cell death threshold, leading to nonspecific assay results. The authors proposed to compare assay activity to baseline toxicity—the nonspecific accumulation of chemicals in cell membranes that disrupt cellular function at critical concentrations. Even without a specific target, compounds can cause potent toxicity due to hydrophobicity, potentially skewing assay results. They found that 37–87% of active hits in the six Tox21 assays were likely due to cytotoxicity burst, while only 2–14% were specific.^12^ This is plausible given assay responses can arise from multiple mechanisms, not all representing compounds with the intended pharmacological action.^13^ Hence, this illustrates that, either directly or indirectly, assay endpoints which one measures (or predicts) can be correlated with confounding factors (often cytotoxicity), and not considering the latter in both measurements and predictive models gives misleading results.

Our originally anecdotal finding that a large proportion of the well-predicted assays in prior benchmarks were cytotoxicity/viability related prompted us to test and discover that simple cell counts are surprisingly predictive in many of these benchmark assay tasks. If simple metrics like cell count are driving much of the predictive power, it calls into question the added value of more complex features, such as Cell Painting or gene expression profiles. We identified three practical implications of this in this work: (1) biased benchmark datasets have a high proportion of cell viability assays; a fairly easy-to-predict assay endpoint that should not be overemphasized in benchmarking studies, (2) many assays in benchmark datasets are specific to targets but the active chemicals in the dataset also impact cell viability and the actual readout is confounded by effects like cytotoxicity burst^12^, and (3) there is an absence of baseline models trained only on cell counts that can demonstrate the comparative advantage of a model trained on more complex phenotypic profiles.

Establishing simple baseline cell count models is essential in machine learning to assess the value of more complex solutions, akin to ‘Occam’s Razor’ when choosing the simplest model to explain a given observation.^14^ Such baselines can include basic molecular features like ‘atom counts’, which have shown strong performance in a virtual screening setting despite lacking structural information^15^, or mean or majority-class classifiers in regression and classification settings, respectively.^1,16,17^ We propose a baseline machine learning model using cell count to assign activity probabilities, as it proves surprisingly effective in many assays, particularly human tumor cell line growth inhibition assays. We recommend that future studies carefully select assay categories for benchmarks and establish baseline cell count models to realistically gauge performance improvements when using high-dimensional phenotypic data, such as images, mRNA, or protein profiles, and present five related recommendations for the machine learning community.

## METHODS

### Cell Painting data

We used the Bray Cell Painting dataset^18^ (cpg0012-wawer-bioactivecompoundprofiling) in this work, which contains images of cells treated with over 30,000 chemical perturbations. We used the dataset processed by Seal et al.^19^, for each plate, where the average feature value of the DMSO controls was subtracted from the average feature value of the perturbations and then divided by the standard deviation for the DMSO samples on the plate. Finally, the median feature value calculated for each compound and dose combination. For replicates, the median feature value was considered only for doses within one standard deviation of the mean dose across all perturbations of the same compound. We use the Cell Painting descriptor “Cells_Number_Object_Number”, which correlates directly with cell count, as the sole feature in our cell-count baseline cell count models.

### Bioactivity data description

We next gathered three annotated assay benchmarking datasets from the literature: Moshkov et al.^9^, Hofmarcher et al.^8^, and Ha et al.^11^ (Table 1). The Moshkov dataset contains bioactivity data from 270 assays, predominantly cell-based and toxicity-related (e.g., growth inhibition, viability) for 16,170 compounds, with 13.4% completeness and 2.7% active data points (defined from individual assay hitcalls). The Hofmarcher dataset, sourced from ChEMBL, comprises data from 209 assays, featuring quantitative pChEMBL data from individual protein targets, as well as tumor cell line growth inhibition assays, for 10,573 compounds, with 2.5% completeness and 34.7% active data points. The Ha dataset, an update on the Hofmarcher dataset, comprises data for 201 assays from ChEMBL, focusing on protein target activity for 10,526 compounds, with 2.6% completeness and 34.7% active data points. These datasets have been used before to evaluate novel machine learning algorithms predicting compound activity using Cell Painting descriptors, making them suitable for comparing our baseline cell count models.^8–11^

**Table 1.** Description of commonly used datasets compiling publicly available annotations for compounds in the Cell Painting. Completeness: the percent of compound-task pairings that have measured activity data; the percentage of active data points reflects the fraction of compound-assay pairings marked as active (bioactive, toxic, etc.).

| Dataset | Number of compounds | Number of bioactivity endpoints ('tasks') | Type of tasks | Completeness | Percentage of active data points |
| --- | --- | --- | --- | --- | --- |
| Moshkov et al.<sup>9</sup> | 16,170 | 270 | Mostly cell-based compound activity and toxicity | 13.4% | 2.7% |
| Hofmarcher et al.<sup>8</sup><br>(also used in Sanchez et al.) | 10,573 | 209 | Protein target/bioactivity and tumor cell line growth inhibition assays | 2.5% | 34.7% |
| Ha et al.<sup>11</sup><br>(subset of Hofmarcher et al.<sup>8</sup>) | 10,526 | 201 | Protein target/bioactivity and tumor cell line growth inhibition assays | 2.6% | 34.7% |
| This Study | 1,349 | 23 | Protein target/bioactivity | 50.8% | 27.2% |

### Comparing active with inactive compounds

Based on binary activity annotations in each of the three datasets, we compared the cell count distributions between active and inactive compounds for each assay using an independent t-test (as implemented in scipy.stats^20^). To mitigate bias from unequal group sizes, we sampled both groups to match the size of the smaller group. We then analyzed the distribution of the Cells_Number_Object_Number feature for active versus inactive compounds.

### Predictivity of cell count

We systematically assessed the predictive utility of the cell count feature across multiple assays by evaluating classification performance at different thresholds. For each assay across all three datasets, binary predictions were generated using a probabilistic scoring approach (logistic regression with fixed weight of -1) based on the Cells_Number_Object_Number feature. A sigmoid-like function (eqn. 1) was applied to transform feature values into probability scores, mapping the difference between each value and a specified threshold to a probability score between 0 and 1.

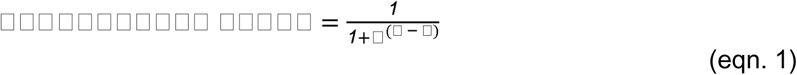

Where value is the raw feature LJ for a specific compound, in this case, the Cells_Number_Object_Number, and LJ is the chosen threshold used to center the transformation.

We then evaluated how varying thresholds on the cell count distributions impacted the ability to predict assay outcomes. The separation of classes was assessed using balanced accuracy at each threshold (probability scores equal to or above 0.5 as positive and those below as negative), emphasizing the ability of the cell count feature to distinguish positive from negative classes. Since our goal was solely to separate classes and not to train and test an ML model, a low balanced accuracy score of 0.10 was interpreted as 0.90, given that the thresholding model is fixed to classify compounds that decrease cell count as active (and often makes opposite predictions). This approach emphasizes our focus on class separation rather than absolute predictions, meaning the assay-specific definition of activity (e.g., whether increased cell counts indicate activity) does not affect our interpretation.

### Predicting compound activity from phenotypic profiles and chemical structures

The Moshkov dataset^9^ uses binary classification across 270 tasks, with scaffold-based splits for training and testing defined by the original authors. We excluded 40 assays, which did not contain enough data points to guarantee that at least one active and one inactive compound is included in the training and test data that Moshkov et al used for each cross-validation fold. A Logistic Regression classifier, using sklearn’s default hyperparameters, was trained using “Cells_Number_Object_Number” as the sole feature to predict each binary endpoint. For comparison, a separate logistic regression model was trained using all CellProfiler features for each endpoint. L2 regularization with C=0.001 and balanced class weighting parameters ensured the model handled the high feature dimensionality and class imbalance well. Five-fold cross-validation was performed using the splits provided by the authors, and performance was measured using the Area Under the Receiver Operating Characteristic Curve (AUC) in combination with the Area Under the Precision-Recall-Gain Curve (AUPRG).^21^ AUPRG was selected as an additional metric due to the large class imbalances in most of the endpoints within the Moshkov dataset, where 232 of the original 270 tasks had at least five times more inactive than active compounds and 185 had at least a 10:1 inactive-to-active ratio. AUPRG provides a universal baseline, which is comparable across assays with different class distributions (unlike Area Under the Precision-Recall Curve), and is less impacted than AUC by the abundance of negative class examples that are present in these classification tasks. Using both metrics, therefore, provides a more comprehensive overview of model performance and one less susceptible to identifying models with low precision as highly effective in this extremely unbalanced data scenario.

### Accurate prediction biological assays with Cell Painting and convolutional networks

To train baseline cell count models, we used the Hofmarcher dataset with the same train-validation-test datasets as in the original publication (70-10-20 splits repeated three times for 209 tasks).^8^ An XGBoost classifier was trained for each assay using only the cell count feature, with hyperparameter optimization performed via randomized search across a predefined grid in a 5-fold stratified cross-validation. The baseline cell count models were trained on the 80% training set and evaluated on the 20% test set. This process was repeated for all three train-test splits provided by the authors. Model performance was assessed using the AUC, allowing for direct comparison with results from Hofmarcher et al.

### Comparison to contrastive learning (CLOOME): bioactivity prediction

We next compared the performance of a logistic regression (LR) model trained on the cell count feature to Sanchez-Fernandez et al.^10^, who trained an LR model on CLOOME embeddings for each of the 209 targets in the Hofmarcher dataset to compare feature spaces like-to-like, while keeping all other parameters identical. Since CLOOME embeddings have access to chemical structures, two baseline cell count models were considered: (1) using cell count as the sole feature and (2) using cell count along with molecular weight and logP (calculated using RDKit) as features. The first baseline excludes any chemical information, while the second baseline, although not explicitly trained on chemical features, incorporates some chemical-related information indirectly through molecular weight and logP. Employing the same 70-10-20 data splits, multiple logistic regression models were trained per assay with varying regularization strengths, selecting the best based on validation AUC. Optimal models were then evaluated on the test set. This process was repeated for all three train-test splits provided by the authors, with evaluations based on the mean AUC for each assay.

### Few-shot learning for molecule activity prediction

We next compared baseline cell count models to those developed by Ha et al., who used 161 assays for training a few-shot learning model and 18 separate assays as the test set.^11^ Since no training data was available for the test assays, the baseline cell count model was trained by combining the outcomes of all 161 training assays into a single endpoint. This approach encourages the model to capture general trends rather than assay-specific information, aligning with our hypothesis that this generic information might primarily reflect cell death. Hyperparameter optimization for the XGBoost model was performed using a 5-fold stratified cross-validation strategy, with cell count as the sole feature. Model performance was evaluated individually on each of the 18 test tasks designated by the authors, using AUC as the performance metric.

### Comparing label-free Cell Painting profiles brightfield images with cell count baseline

We next used the test dataset provided in Cross-Zamirski et al^22^ which included 77 negative DMSO wells, 26 positive wells, and 170 compound perturbation wells. We used 4 feature spaces, namely (a) Cell Painting profiles from the ground truth, (b) features from predicted images from the conditional Wasserstein Generative Adversarial Network with gradient penalty (cWGAN-GP) model, (c) features from predicted images from the U-Net model as released by the authors, and (d) the baseline cell count model using the Cells_Neighbors_SecondClosestObjectNumber_5 feature from the ground truth profiles. To identify compounds with phenotypic similarity to positive controls, we first used a k-Nearest Neighbors (k-NN) classifier (k=5) with Euclidean distance in each of the above feature spaces to distinguish between the positive and negative control. The model was trained and evaluated through 100 iterations, where the training data was randomly resampled in each iteration to ensure balanced representation of both classes. Predictions for each of the 170 compound perturbation wells were aggregated using majority voting across all iterations, identifying those test compounds predicted as the positive control.

### Comparing feature extraction from high-content imaging screens with cell count baselines

We next used the dataset released by Comolet et al.^23^ which consisted of fibroblasts from 20 donors treated with three compounds (CP21R7, Pemigatinib, and Y-39983-HCl) at two concentrations (0.2 and 1.0 μM) each in order to compare the ability of various Cell Painting feature space to identify compounds from controls. The authors performed confounder-correction (plates, wells, rows, columns, donor), normalized between 0 and 1, and averaged at the well level to mitigate noise and emphasize the predominant effects observed across the majority of the cell population. ^23^ We used two features spaces to compare the ability to separate perturbations from DMSO controls–first, the ScaleFEx’s features which included all extracted morphological and phenotypic descriptors from cell imaging assays and second, only the MitoCount feature extracted by ScaleFEx, which was the closest related feature to cell count. We reproduced the methods described by Comolet et al to assess the accuracy of a logistic regression (LR) model designed to distinguish each drug from the control (DMSO). For each drug and concentration, a balanced training set was created by sampling DMSO controls to match the minority class. For models using ScaleFEx features, a Recursive Feature Elimination (RFE) and logistic regression model was employed to identify the most predictive, non-redundant features for classification. For the baseline cell count model using MitoCount, no feature selection was used. A logistic regression model was trained and validated using leave-one-plate-out cross-validation (which results in information leak of compounds which was spread across all plates), that is, in each fold, training is performed on all but one plate, which is held out for testing. The models are trained iteratively to classify wells treated with each drug and DMSO controls. The values were corrected for plate, donor, column, and row effects and normalized to a range of 0 to 1. A Mann-Whitney U test was used to assess statistical differences between a drug and DMSO control, with Bonferroni adjustment applied to account for multiple comparisons.

### Evaluating the effects of concentration-dose response in Cell Painting

We hypothesized that the signal in Cell Painting depends on compound concentration and that single concentrations in public datasets may be suboptimal. As an illustrative example, we generated dose-response Cell Painting data for six compounds: aminohexylgeldanamycin (Hsp90 inhibitor), givinostat (HDAC inhibitor), repotrectinib (tyrosine kinase inhibitor), I-BET282 (pan-inhibitor of all eight BET bromodomains), bisindolylmaleimide V (protein kinase C inhibitor), and PP58 (Src inhibitor), in 2 replicates and ten doses.

### Benchmarking Cell Painting for curated Protein-Target data

To evaluate the use Cell Painting assay to predict protein targets of compounds, we curated a dataset from PubChem, ChEMBL, and DrugMatrix based on compounds for which Cell Painting profiles were provided by Bray et al. Assays were filtered to include activity types IC50, EC50, AC50, GI50, and Ki for single proteins in *Homo sapiens* (non-mutant and non-recombinant proteins), with “=” as activity relation. To harmonize activity annotations, the ChEMBL-assigned activity_comment field was categorized into predefined functional classes, including “inhibitor,” “active,” “inactive,” “substrate,” “antagonist,” “inverse agonist,” and “agonist,” and these classifications were compared with pChEMBL values, where compounds with pChEMBL values > 6.5 were considered active. Entries with ambiguous annotations or missing data were removed. Compounds with low cell count in the Cell Painting assay were excluded, which was defined as the mean cell count minus two standard deviations. Assays with insufficient class diversity (<20 compounds per class) were removed. These filtering steps resulted in a high-quality dataset comprising 23 protein-target biological assays with binary activity labels (active/inactive) for a total of 1,349 compounds.

We evaluated the predictive performance of two feature sets to predict protein targets: the full Cell Painting profiles extracted using CellProfiler and a baseline cell count model based on cell count. The baseline cell count model used a single morphological feature representing cell count, while the Cell Painting feature set encompassed diverse cellular morphology descriptors. To address class imbalance, class weights were calculated using the balanced strategy (classes with fewer samples receive higher weights), ensuring equal contribution of positive and negative classes during model training. Models were trained using stratified 3-fold nested cross-validation to maintain balanced class distributions in both training and test sets. Logistic Regression was employed for the baseline cell count model with the cell count feature, while Random Forest Classifiers were used for the multi-dimensional Cell Painting features. For hyperparameter optimization, 2/3rd of the data was used in the inner cross-validation loop, applying RandomizedSearchCV to tune parameters. This included regularization strength for Logistic Regression and tree depth, leaf size, and the number of estimators for Random Forests. The best-performing models were selected based on AUC-ROC scores from the inner cross-validation loop. Classification thresholds were optimized using Cohen’s Kappa, evaluated across a range of probability thresholds to maximize agreement between predicted and true labels in the training data. Final model performance was assessed on the outer test set using metrics such as Balanced Accuracy, AUC-ROC, AUC-PR, Matthews Correlation Coefficient (MCC), and Cohen’s Kappa. To ensure robustness, random label scrambling was performed for each assay, and models were retrained on the scrambled labels to confirm that their performance was not driven by chance.^24^

### Statistics and Reproducibility

All statistical analyses were performed as implemented in scikit-learn.^25^ All code is available at https://github.com/srijitseal/The_Seal_Files. All data is released via https://doi.org/10.5281/zenodo.14838604.

## RESULTS AND DISCUSSION

### Multiple bioactivity tasks are highly correlated with the simple cell count feature

We first evaluated baseline cell count models trained on a single image-based feature, namely cell count based on segmentation of nuclei in a DNA-stained image, across four previously published bioactivity prediction benchmarks: (a) Moshkov et al.^9^, (b) Hofmarcher et al.^8^, (c) Sanchez-Fernandez et al. (CLOOME)^10^, and (d) Ha et al.^11^ We began by assessing the relationship between bioactivity prediction tasks and changes in cell count by comparing cell count distributions for active and inactive compounds across three benchmark datasets. Significant differences between both sets were observed in all datasets (Figure 1 a-c) when using an independent samples t-test (p <= 1.00e-04), suggesting that cell count alone might contain substantial predictive signal for many biological assays.

**Figure 1.**
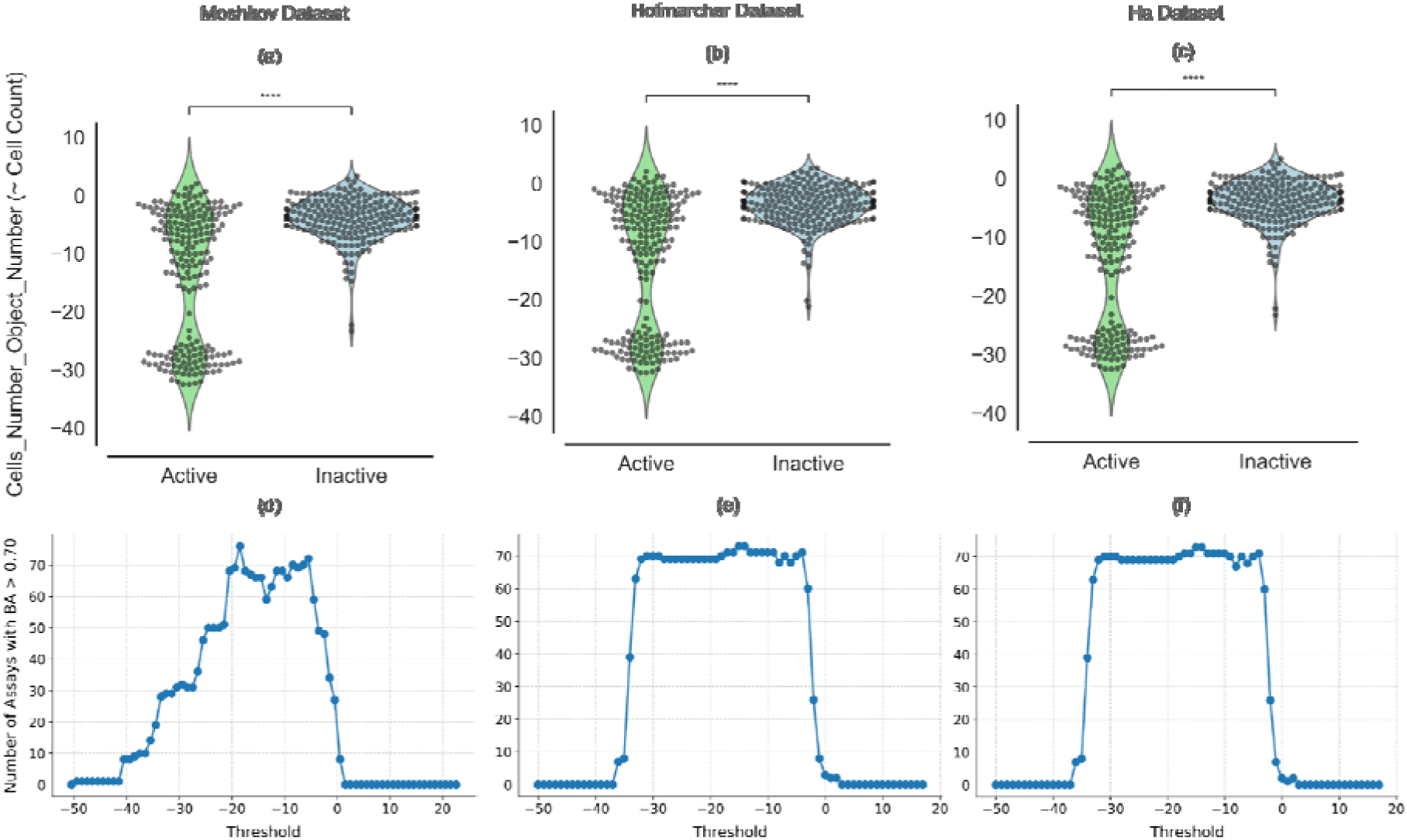
Comparison of the distribution of the cell count feature (“Cells_Number_Object_Number”) across (a) 270 assays in the Moshkov et al.^9^, (b) 209 assays in the Hofmarcher et al.^8^ (which was also analyzed in Sanchez-Fernandez et al.^10^), and (c) 201 assays in the Ha et al.^11^, split by active and inactive compounds and balanced to the same number of active and inactive compounds per assay. ****: p <= 1.00e-04 t-test independent samples with Bonferroni correction. Number of assays with balanced accuracies>0.70 when predicting assay outcomes in the (d) Moshkov dataset, (e) Hofmarcher dataset, and the (f) Ha dataset, by thresholding the normalized cell count feature at various cutodds.

We further analyzed the predictivity of the cell count feature for benchmark datasets used in prior studies and found that solely applying a range of thresholds of cell counts successfully predicted 32% of assays (66 out of 209) in the Hofmarcher dataset, with mean balanced accuracy over 0.70 (Figure 1d), with 64 of the 66 assays being human tumor cell line growth inhibition assays. Thresholds of cell counts successfully predicted 22% of the tasks in the Ha dataset and 28% in the Moshkov dataset (Figure 1e and 1f). This indicates that these datasets contain a substantial number of assays whose outcomes primarily correlate with cell count and therefore are not ideally suited for assessing data modality or model effectiveness across a broad spectrum of assays.

### Cell count matches performance of models trained on Cell Painting and gene expression profiles on existing benchmarks

We next investigated the information content in the cell count feature; we followed the same train-test splits for each of the studies reanalyzed in this work, and compared our baseline cell count model performance to those models trained on the higher dimensionality data of Cell Painting and gene expression profiles.

In the Hofmarcher dataset^8^, which was also analyzed in Sanchez-Fernandez et al.^10^ and of which the Ha dataset^11^ is a variation, the baseline cell count models (mean AUC = 0.68L±L0.21) achieved similar performance compared to the supervised fully-connected neural network (FNN) model based on the full Cell Painting profiles (mean AUC = 0.67L±L0.19), while the convolutional neural network architectures of ResNet model trained on Cell Painting images directly (mean AUC = 0.73L±L0.19) performed better (Table 2, rows e,f,g). We observed a significant overlap of assays well-predicted across all three models (66 were well-predicted, AUC>0.8), with the two more sophisticated image-based models collectively predicting only 19 additional assays over the baseline cell count model (Figure 2a). We found the performance of all models, whether using cell count or Cell Painting features, were largely based on human tumor cell line growth inhibition assays (Figure 2d). Ideally, benchmarks should not over-represent a single type of assay, especially viability assays, given that it is a readout that can be well-predicted by a single feature such as cell count (albeit with differences due to assay conditions such as cell type and timepoint).

**Table 2.**
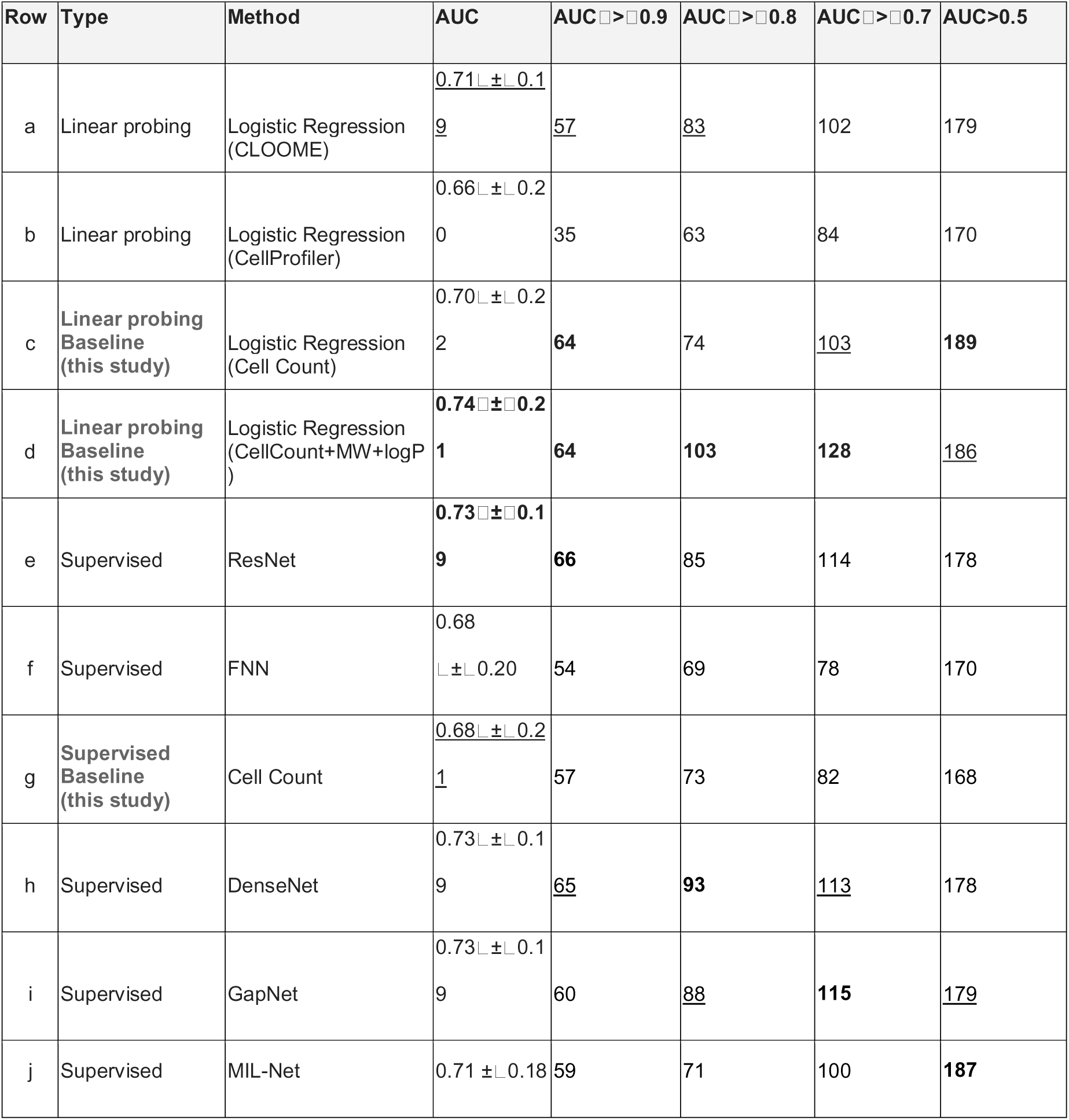
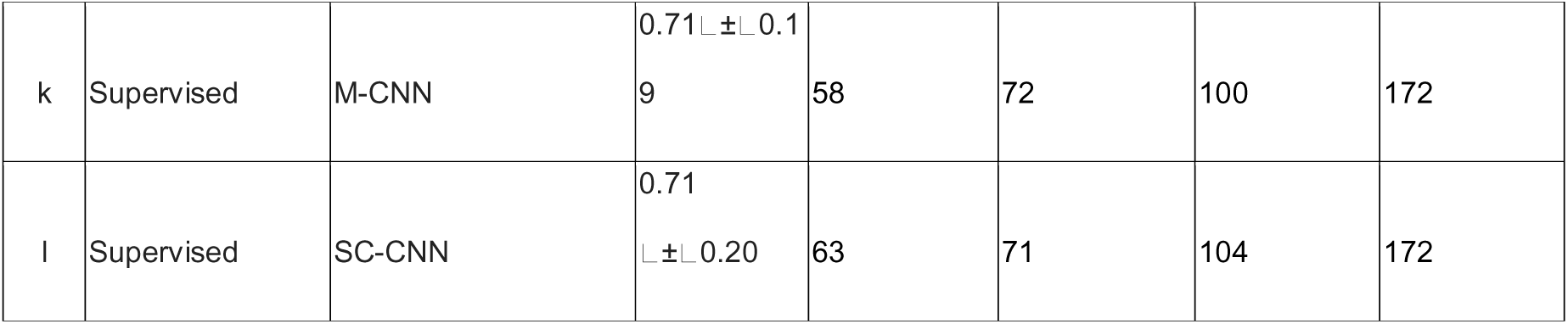
Performance metrics across 209 prediction tasks in the Hofmarcher dataset. These baseline cell count models are compared to models used in Hofmarcher et al.^8^ and Sanchez-Fernandez et al.^10^ (based on the complete Cell Painting features). Also visualized in Figure 3. Bold and underlined values indicate best and second-best performance.

**Figure 2.**
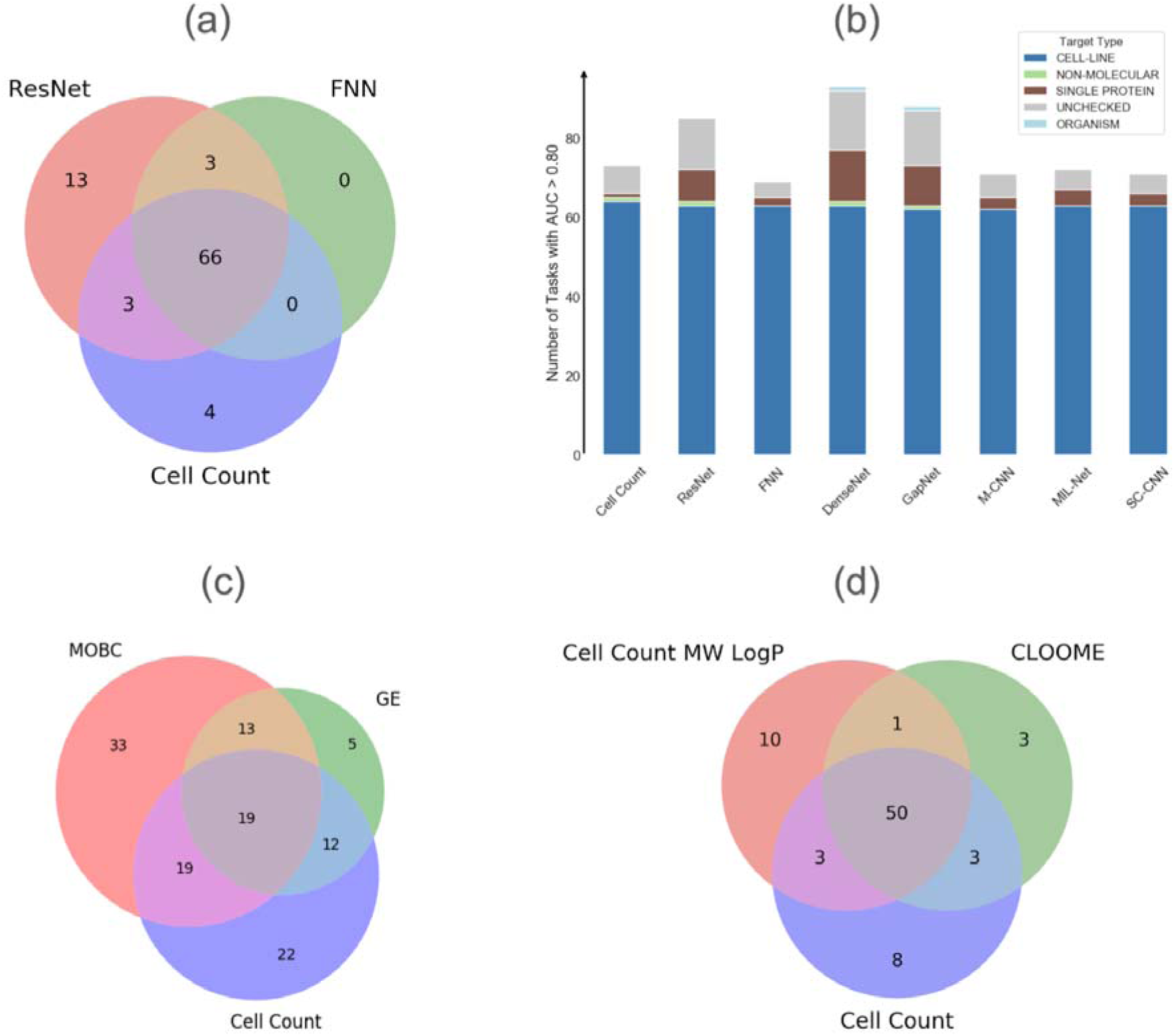
The number of assays predicted with (a) AUC > 0.8 and (b) assay types predicted with AUC > 0.8 in the Hofmarcher dataset, (c) the number of assays with AUC > 0.7 in the Moshkov dataset, and (d) number of assays with AUC > 0.9 with linear probing models using CLOOME image and chemical structure-based embeddings compared to baseline cell count models are compared to models used in Sanchez-Fernandez et al.^10^ Multi-task neural network (results from Moshkov et al.), trained using either CellProfiler morphological profiles which have been batch-corrected (MOBC), or gene expression data (GE). Baseline cell count models were trained only using the “Cells_Number_Object_Number” morphology feature (cell count). Differing AUC cutoffs are used to be consistent with the originally published results for each dataset. FNN: Fully connected neural network, ResNet: residual neural network, CLOOME: Contrastive Learning and leave-One-Out-boost for Molecule Encoders.

**Figure 3.**
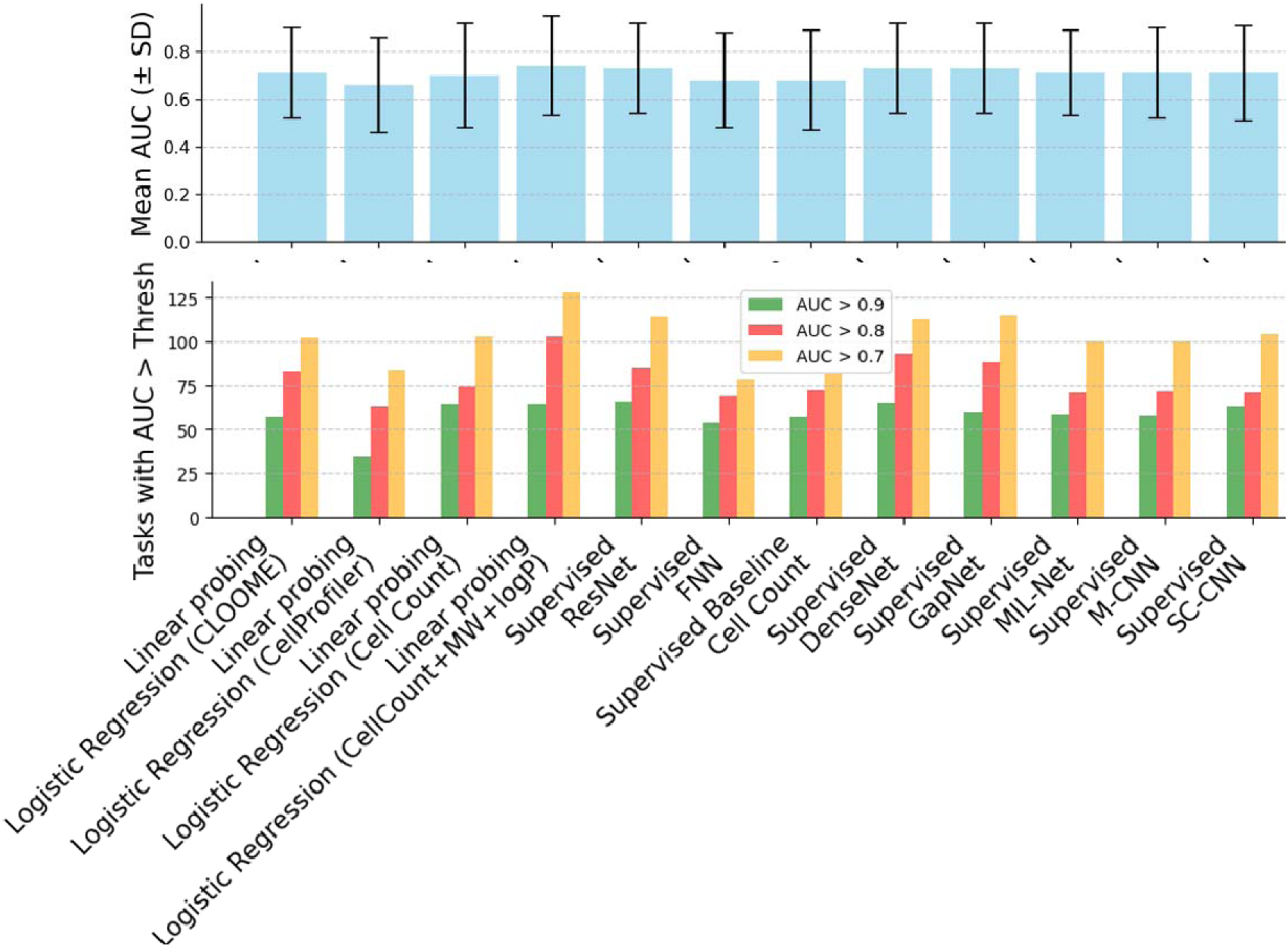
Top: Mean AUC values (± SD) for each model, categorized by method and type (Linear Probing, Supervised, Baseline, Control) for the tasks in the Hofmarcher dataset. Error bars represent the standard deviation for each model’s AUC across experiments. Bottom: Assays where the models achieved AUC greater than the chosen thresholds. Bars represent non-cumulative counts. Y-scrambling control demonstrates minimal performance as a control.

The Hofmarcher dataset contains cell-based assays as well as biochemical binding assays using purified protein targets. Out of the 66 unique protein targets in the Hofmarcher dataset, cell count predicted only one protein target at a performance level of AUC>0.8, while the convolutional neural network architectures of ResNet and DenseNet using images predict six targets and eleven targets respectively (Figure 2b). This shows that while most methods perform similarly to baseline cell count models when evaluated on all assay types, when predicting small molecule activity on protein targets, Cell Painting images achieve high predictivity for many unique protein targets (for example, ATAD5, IDH1, ATXN2, MAPT, SMAD3, Hsf1, VDR, RAPGEF3, BRCA1, TDP1, and GMNN for convolutional neural network DenseNet model), many of which relate to DNA damage. This shows that the Cell Painting images are able to predict assay outcomes from many unique protein targets where the baseline cell count model fails.

We next investigated whether this substantial overlap in predictable assays was only feasible because of Cell Painting images’ inherent ability to count cells or whether other unbiased profiling methods, such as mRNA profiling, might capture cytotoxicity/viability equally well. Using the Moshkov dataset^9^, we found a substantial intersection between assays predicted by the baseline cell count model and Moshov’s multi-task neural network trained on gene expression data (Figure 2c), a similar result as for models using Cell Painting data. 31 out of 49 (63%) assay endpoints well-predicted using gene expression profiles (AUC > 0.7, threshold as used by the authors of the study, see Table 3d) were predicted equally well (AUC > 0.7) by using the baseline cell count model. Only five assays were uniquely well-predicted by gene expression profiles, while in contrast, 33 assays were uniquely well-predicted by incorporating the entire Cell Painting feature set and 22 assays were uniquely predicted by the baseline cell count model. Overall, we found that the performance of the baseline cell count models using a single cell count feature could capture much of the information contained in Cell Painting and gene expression data for the assays evaluated in prior studies.

**Table 3.** Performance metrics across 230* prediction tasks in the Moshkov dataset when using a baseline cell count model, compared to late fusion models evaluated in Moshkov et al.^9^ Bold and underlined values indicate best and second-best performance. CS: chemical structure, GE: gene expression, MOBC: morphology, batch corrected - Cell Painting.

| Row | Method | Features | Mean AUC | AUC $\geq 0.9$ | AUC $\geq 0.7$ | AUC > 0.5 |
| --- | --- | --- | --- | --- | --- | --- |
| a | Logistic Regression Baseline | Cell Count | <u>0.611</u> $\pm 0.145$ | 11 | 72 | 179 |
| b | Logistic Regression | MOBC | $0.613 \pm 0.156$ | **13** | 83 | 179 |
| c | Multi-task Neural Network<sup>9</sup> | MOBC | $0.628 \pm 0.170$ | 9 | 84 | 189 |
| d | Multi-task Neural Network<sup>9</sup> | GE | $0.596 \pm 0.140$ | 5 | 49 | 180 |
| e | Multi-task Neural Network (Late Fusion)<sup>9</sup> | CS, MOBC | $0.649 \pm 0.183$ | **13** | **99** | 190 |
| f | Multi-task Neural Network (Late Fusion)<sup>9</sup> | GE, MOBC | <u>$0.650 \pm 0.149$</u> | <u>10</u> | 89 | **196** |
| g | Multi-task Neural Network (Late Fusion)<sup>9</sup> | CS, GE, MOBC | **$0.658 \pm 0.159$** | <u>10</u> | <u>94</u> | <u>195</u> |
| h | y-Scrambling Control | MOBC | $0.483 \pm 0.207$ | 1 | 15 | 97 |

*\*40 assays have been excluded, which did not contain enough data points to guarantee that at least one active and one inactive compound is included in the training and test data for each fold in cross-validation. Therefore, results from Moshkov et al.<sup>9</sup> have been updated to exclude these assays.*

This analysis highlights the value of using cell counting models as an initial, interpretable approach before employing more complex features. In this manner, endpoints that can be easily predicted can be excluded for further in-depth analysis, or at least overall results can be put into context by comparing how much value each modality adds on top of using cell count.

### Baseline cell count models show comparable AUC to meta-learning approaches

We next compared our baseline cell count models with the single-task, multi-task, and meta-learning models using Cell Painting data used by Ha et al.^11^ We found that the baseline cell count model achieved a median AUC of 0.59 on this benchmark, whereas the “protonetcp+” model, a metric-based meta-learning model using embeddings from Cell Painting features, attained an AUC of 0.64 (Table 4). Notably, with regards to the exceptional performance on specific assays, such as CHEMBL2114784^26^ (AUC of 0.87), a viability screen for the primary quantitative high-throughput screening (qHTS) aimed at identifying inhibitors of ATXN expression, the baseline cell count model achieved an AUC of 0.82. The similarity in AUC scores suggests that the baseline is competitive with the more complex model, likely because the assay is linked to cell viability.

**Table 4.** Comparison of Protonet CP+ model (at support set size 64) with baseline cell count model as benchmarked on FSL-CP in Ha et al.^11^

| ChEMBL ID | Protonet CP+ (64) AUC | Baseline Cell Count model AUC | Baseline Cell Count Model with labels reversed AUC |
| --- | --- | --- | --- |
| CHEMBL1614530 | 0.65 | 0.60 | 0.40 |
| CHEMBL1040692 | 0.61 | 0.53 | 0.47 |
| CHEMBL1741325 | 0.68 | 0.54 | 0.46 |
| CHEMBL1741323 | 0.7 | 0.56 | 0.44 |
| CHEMBL1741324 | 0.59 | 0.57 | 0.43 |
| CHEMBL3562136 | 0.74 | 0.75 | 0.25 |
| CHEMBL1738598 | 0.54 | 0.43 | 0.57 |
| CHEMBL1738606 | 0.63 | 0.57 | 0.43 |
| CHEMBL1738312 | 0.57 | 0.65 | 0.35 |
| CHEMBL1794311 | 0.77 | 0.62 | 0.38 |
| CHEMBL1794486 | 0.63 | 0.60 | 0.40 |
| CHEMBL1794499 | 0.6 | 0.70 | 0.30 |
| CHEMBL1794584 | 0.8 | 0.75 | 0.25 |
| CHEMBL1964081 | 0.69 | 0.64 | 0.36 |
| **CHEMBL2114784** | **0.87** | **0.82** | 0.18 |
| **CHEMBL2114807** | **0.77** | **0.24** | **0.76** |
| CHEMBL2354287 | 0.54 | 0.50 | 0.50 |
| CHEMBL2114817 | 0.61 | 0.50 | 0.50 |

Furthermore, Ha et al. posited that pre-training significantly enhanced model performance on other assays, such as CHEMBL2114807.^11,27^ On further examination, we found this qHTS assay as designed to identify small molecule activators of BRCA1 expression. The labeling for this assay was as follows: compounds with lower pChEMBL values were classified as activators and labeled ‘1’. In comparison, those with higher pChEMBL values were labeled ‘0’, representing likely inhibitors that could impair DNA repair mechanisms and induce cell death.^27^ By reversing the output of the baseline cell count model to predict low cell counts as ‘0’, we achieved an AUC of 0.76, closely matching the best pre-trained models reported by the authors (AUC = 0.77). Hence we can conclude that it is key to use data in a suitable way when training models and also that cell counts are very predictive for this particular endpoint.

These results underscore a cautionary reminder regarding interpreting AUC values: models predicting assays related to cytotoxicity cannot be easily compared with each other; often, in the case of meta-learning, test assays predict the opposite outcome of the training assays and a model, being unaware of this, predicts precisely the opposite of the assay outcomes. Given that the user always knows what assays it intends to predict, this task does not need meta-learning but merely reversing the prediction outcomes, as demonstrated here. Thus, AUC values provide only part of, but not the most important part of, performance.^28^

### Contrastive learned features do not outperform cell count features

We next evaluated whether contrastive-learned morphological features learn more than just cell count, in the context of assay prediction. We used features learned using CLOOME (a contrastive learning framework that pairs microscopy images and chemical structures to enhance molecular representation learning) to two logistic regression models: one trained with the cell count feature and another with cell count, and two readily-calculated features of chemical structure: molecular weight and logP.^10^ On the Hofmarcher dataset and the publicly available CLOOME embeddings, we tested linear probing, which is a logistic regression task commonly used to assess the quality of representations learned through contrastive learning. It achieved an average AUC of 0.71 ± 0.19 and 57 well-predicted assays compared to using the cell count feature, which achieved a similar average AUC of 0.70 ± 0.22 and 64 well-predicted assays (Table 2, rows a and b). We found that furthermore the logistic regression model trained on cell count, molecular weight, and logP values achieved an average AUC (0.74 ± 0.21) and 64 assays with AUCL>L0.9, thus even slightly superior to CLOOME (Figure 2d). Overall, this highlights the importance of cell counting as a baseline cell count model and the choice of a suitable task/endpoint for model benchmarking, given there was no observed step-up in performance when using contrastive learning embeddings.

### Appropriate evaluation metrics and assay endpoints are essential when modelling large-scale benchmark datasets

The multi-task neural network models trained on complete Cell Painting profiles in Moshkov et al. achieved extremely high AUC values (1.0) for nine out of the original 270 assays.^9^ Eight of these endpoints contained only one active compound, while one contained two actives. Each endpoint also contained fewer than 52 data points in total. This lack of data, combined with the scarcity of active compounds, made high AUCs achievable by chance. On inspection, nine of the ten unique compounds labelled as active for these assays were cytotoxic^29–31^, leading to significant cell death and low cell density in images (some examples in Figure 4). Therefore, the high performance could be attributed to models effectively predicting cytotoxicity or the fact that test splits in multi-task models often contain very few compounds in the test dataset, rendering the correct prediction of a single active compound by chance plausible.

**Figure 4.**
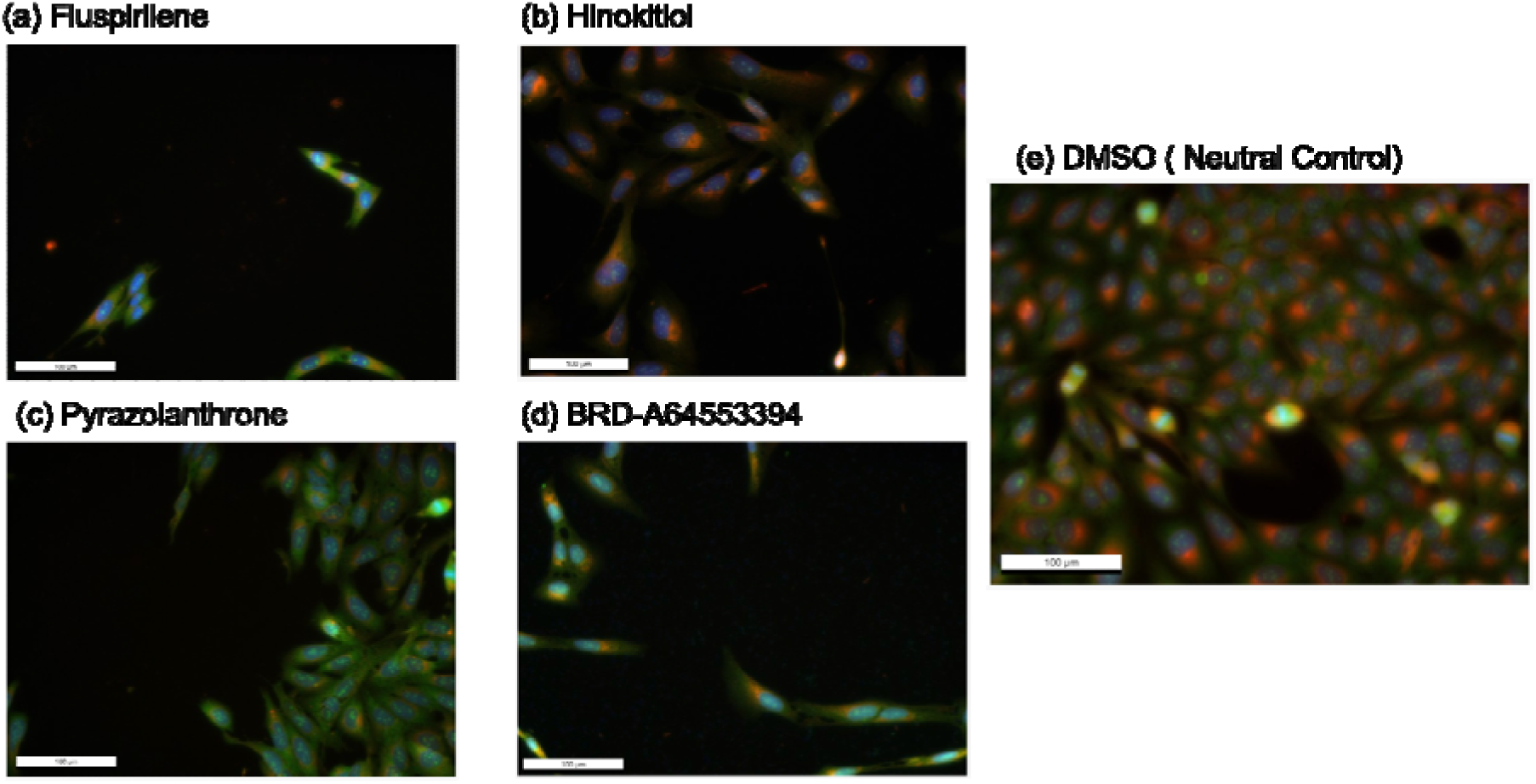
Example images of cells, treated with four different compounds (fluspirilene, hinokitiol, pyrazolanthrone, BRD-A64553394) and neutral control DMSO show significant cell death, dark areas, and low cell density, indicating cytotoxicity (the most extreme cases are bordered yellow). These images (a-d) may not have been used in generating profiles, as a QC pipeline in CellProfiler is likely to reject them and only keep some images with some surviving cells^32^; nevertheless, the images show that these compounds are cytotoxic.

This hypothesis is supported by comparing the AUC scores of the different model approaches after separating the remaining 230 assay endpoints into two groups; 185 assays containing five or more active compounds and assays containing fewer than five active compounds (Figure 5a and 5b). This reveals that a greater proportion of endpoints are predicted with higher AUC when there are fewer active compounds in the data. This is particularly true in the case of our baseline cell count model, where the use of one feature and a simpler model architecture likely leads to less overfitting. Conversely, as the size of the training data increases, a cell count baseline underperforms the two approaches trained using all CellProfiler features. This demonstrates that, for appropriate endpoints, more complex approaches are deriving additional value from high dimensional feature sets beyond cell count.

**Figure 5.**
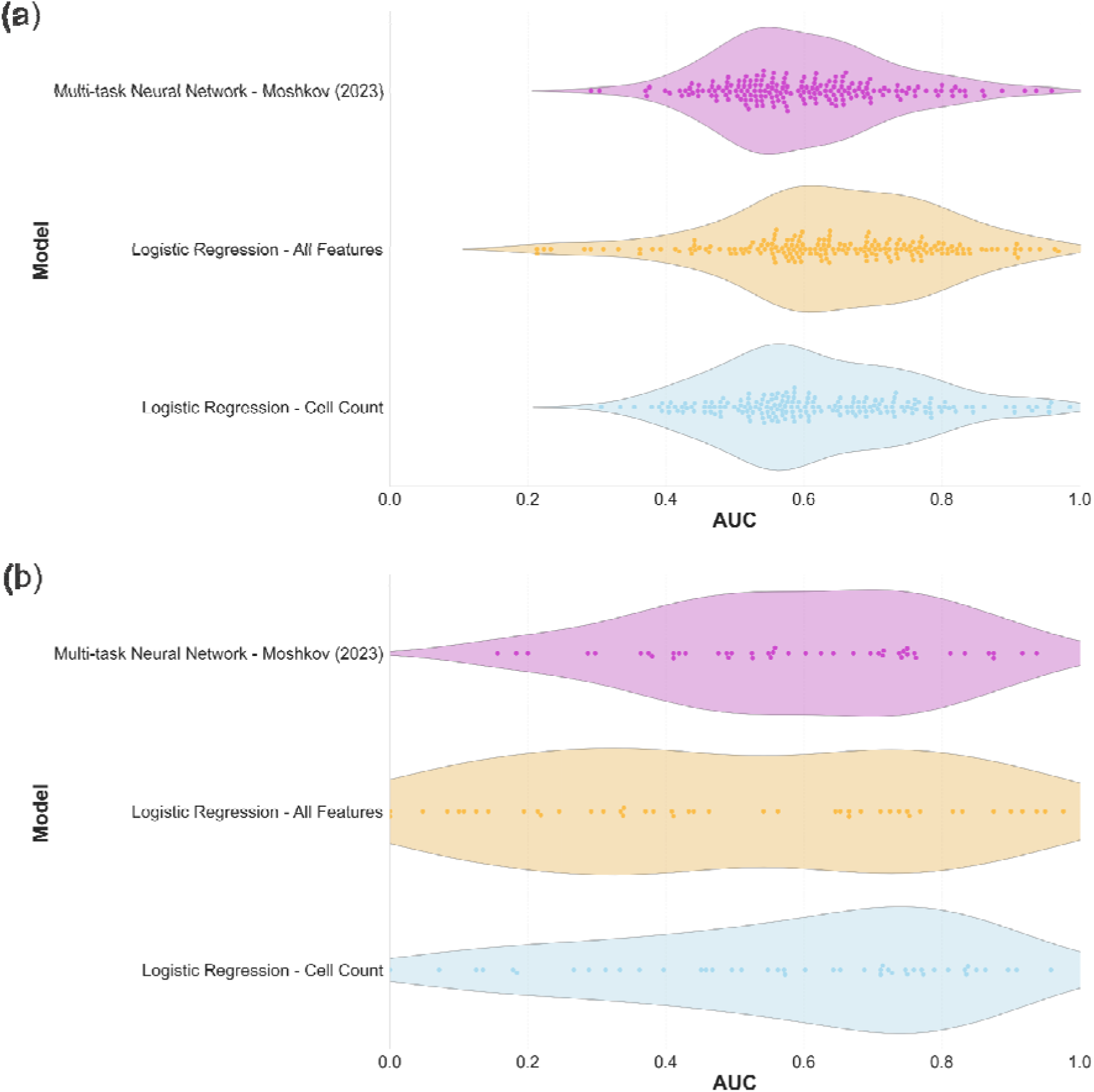
Swamplot showing the distribution of performance of three different models across the assays (dots) when; (a) there are >= five active compounds, (b) < five active compounds in the tested assay. The logistic regression models perform comparably to the more sophisticated multi-task neural network in both scenarios, but (b) demonstrates that, in the case of assays with limited active compounds, using cell count only can lead to superior prediction.

To investigate whether the sole use of AUC could be misleading in terms of judging model effectiveness, for the remaining 230 assays in the Moshkov dataset we plotted the AUC of each model against AUPRG, which as mentioned above provides a universal baseline for comparison between assays which is less impacted by severe class imbalance (Figure 6a and 6b). Thresholds to categorise assay endpoints as “well-predicted” were set at 0.7 AUC and 0.3 AUPRG. Using the more stringent performance criteria meant that just 4 assays were considered to be well-predicted by our logistic regression model (Figure 6a), in contrast to the 83 assays predicted with AUC > 0.7 or 13 with AUC > 0.9 (Table 3b). This reduction shows that AUC alone was insufficient to assess the number of “well predicted” assays.

**Figure 6.**
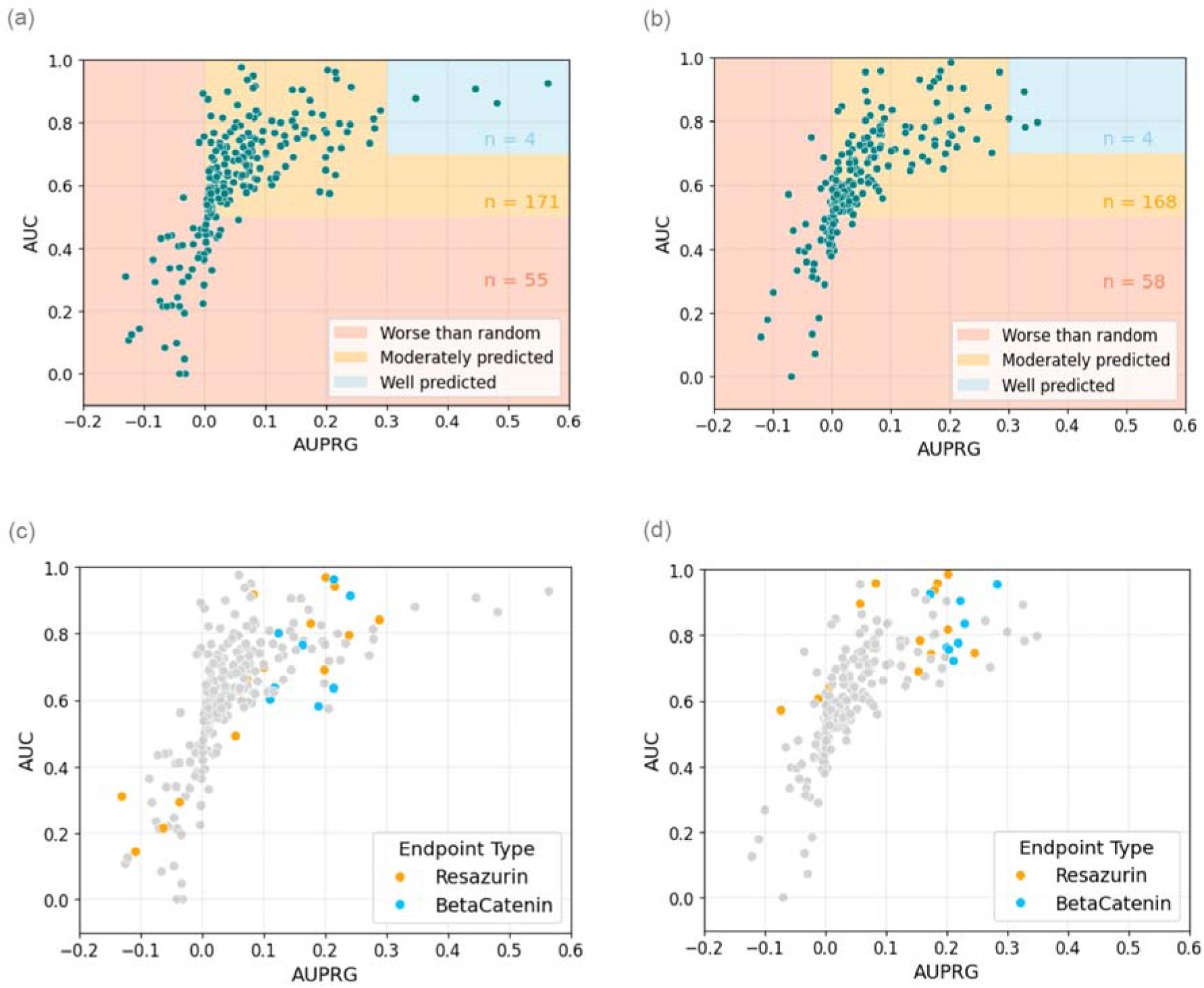
Comparison of the performance of logistic regression models incorporating either; (a) all CellProfiler features (b) our cell count baseline using only the “Cells_Number_Object_Number” feature. Area under the receiver operating curve (AUC) has been plotted against area under the precision-recall-gain curve (AURPG) to appropriately compare performance in the Moshkov dataset where there is significant data imbalance. Specific endpoints of interest have been highlighted relating to either Resazurin (orange) or BetaCatenin (blue) assays for the logistic regression models trained with all CellProfiler features (c) or only the cell count feature (d). These two assays typically identify active compounds that cause reduced cell viability and it is shown above that in these instances the cell count baseline approach (d) demonstrates strong predictive ability.

We also hypothesized that a number of the assay endpoints were highly correlated with cell death, and thus would be predicted with high performance by a baseline model using the cell count feature alone. We therefore focused on two distinct endpoint types within the data; resazurin and Wnt/β-catenin signalling pathway assays. The resazurin assay is commonly used to assess both bacterial and eukaryotic cell viability^33,34^, whilst the Wnt/β-catenin signalling pathway is crucial for cell proliferation and survival.^35^ The Cell Painting experiments used U2OS osteosarcoma cells where Wnt signalling is often active^36,37^; therefore it is likely that active inhibitory compounds would result in reduced cell proliferation, as well as increased apoptosis and cell cycle disruption. These endpoint types accounted for 14 and eight assays, respectively, out of the 230 assays in the data.

Generally, these two endpoint types were well-predicted by our baseline cell count model, with an average 0.806 AUC and 0.154 AURPG, outperforming the 0.70 average AUC achieved by Moshkov et al.’s late fusion of three different data modalities for those endpoints. Only two out of 22 assays (9%) were predicted worse than random by the cell count baseline (Figure 6d). However, the logistic regression model trained on all CellProfiler features actually results in worse model performance on these endpoint types (Figure 6c), with five resazurin assay endpoints predicted with AUC < 0.5. Instead, other assays are predicted with greater AUC/ AURPG scores in their place, resulting in a similar number of assays above our determined performance thresholds (see Table 2a and 3b). This finding supports the necessity to filter these benchmark datasets, to focus on endpoints where applying advanced deep learning techniques will yield real benefits over and above the information contained in cell count.

### Comparing compounds to positive controls is not a good benchmark for ML models

Cell Painting experiments use positive controls that are typically compounds that exhibit a strong phenotypic change, often including a reduction in cell counts. Studies that compare methods by evaluating whether models can predict if a compound is similar to positive controls are likely confounded by cell count differences. For example, Cross-Zamirski et al. demonstrated label-free prediction of Cell Painting profiles from brightfield images.^22^ This approach enabled the clustering of promising compounds with a positive control compound using a k-NN classifier applied to features from actual Cell Painting fluorescent images and predicted ones derived from brightfield images. The authors reported a sensitivity of 62.5% and a specificity of 99.3% when predicting cytotoxicity, with correct classifications for eight toxic compounds.

Among the label-free models, the feature most strongly correlated with the ground truth was Cells_Neighbors_SecondClosestObjectNumber_5 (Pearson correlation = 0.98), which is closely linked to cell count. Using this feature, we evaluated the 170 treatment wells in the test set and identified eight treatment compounds within 1.5 standard deviations of the positive control’s feature value (Figure 7a). When k-NN models were trained on either the full Cell Painting profiles (ground truth) or the single cell neighbor feature, they identified 7 and 19 treatment wells, respectively, as similar to the positive control. In contrast, label-free Cell Painting profiles generated using cWGAN-GP and U-Net models identified 10 and 14 wells, respectively. As shown in Figure 7b, the majority of wells identified by the label-free models were encompassed within the predictions of the baseline cell count model. Because the dataset used by Cross-Zamirski et al. is not publicly shared, it remains unclear whether these well-level predictions align with their reported compound-level classifications. However, the strong correlation observed between the ground truth and the label-free cell count feature suggests that the predicted compounds from both datasets likely overlap. This evidence highlights the importance of including a baseline cell count model when presenting more complex models. Both Cell Painting and label-free models consistently identified compounds causing significant decreases in cell count, similar to the positive control compound mitoxantrone, a known cytotoxicant. Therefore, baseline cell count models serve as a reference point for evaluating and interpreting results from more complex models, particularly for tasks inherently linked to positive controls.

**Figure 7.**
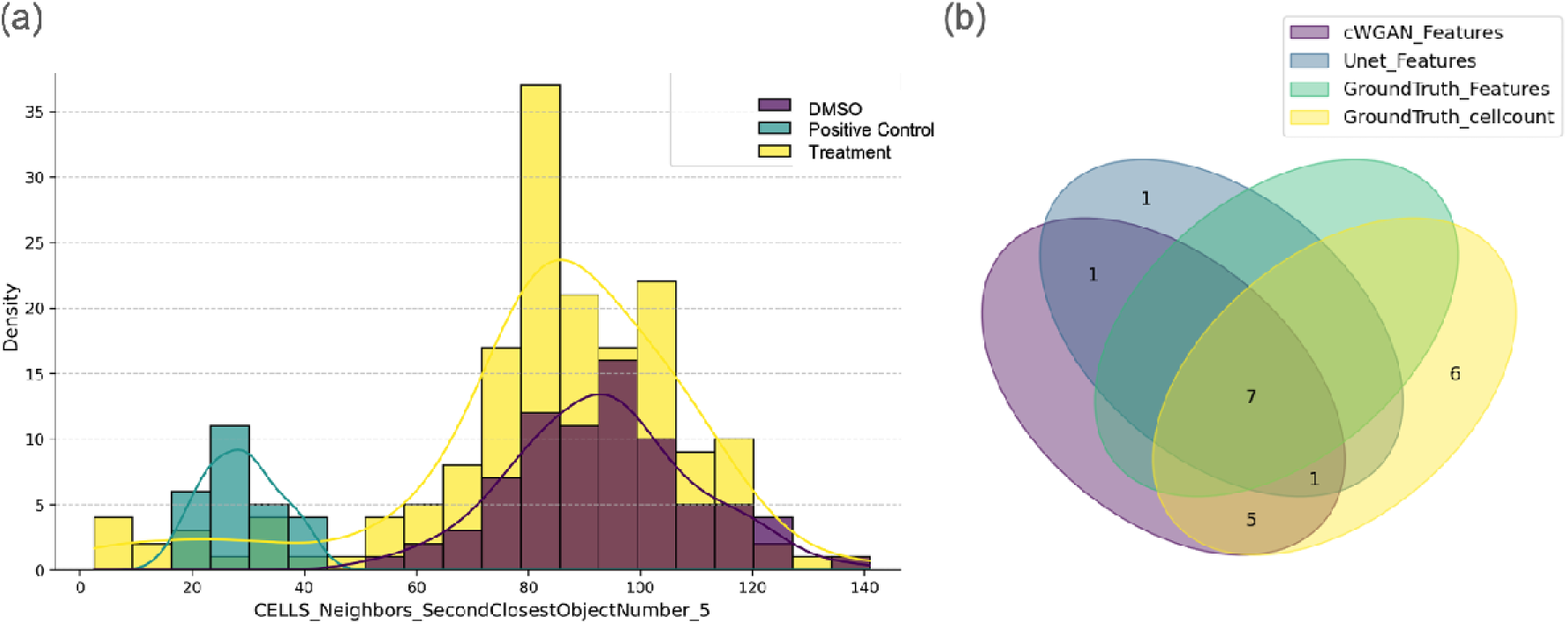
(a) Distribution of the Cells_Neighbors_SecondClosestObjectNumber_5 feature for positive and negative controls, and treatment wells. It shows that positive controls have reduced cell count compared to DMSO controls and most other treatments. (b) Overlap of the wells identified as similar to the positive control by different approaches: the label-free models (cWGAN-GP and U-Net), the full Cell Painting profile and the baseline model using only the cell neighbor feature which captures cell count indirectly.

### Concentration-dependent effects enhance information content in Cell Painting data

Most studies use data from two large public Cell Painting datasets: Bray et al.^18^ and JUMP-CP^38^. These datasets primarily consist of single-dose experiments, where each compound perturbation was tested in multiple replicates but at one concentration, usually 10 μM. A single-dose setup makes it difficult to ascertain whether the compound concentration is optimal to capture unique effects of the compound, or alternatively whether it is higher than desired and begins to capture effects associated with different mechanisms, including those that are outright cytotoxic (conditions where baseline cell count models perform well). Hence, we next explored whether the information content in Cell Painting profiles is concentration-dependent, using datasets with multiple compound concentrations.

To this end, we used data from a recent study by Comolet et al. which developed ScaleFEx, a memory-efficient and open-source pipeline for extracting interpretable cellular features from large high-content imaging datasets. We evaluated the ScaleFEx feature space to identify phenotypic shifts in drug-treated cells. For example, the study validated ScaleFEx features on fibroblasts treated with a single drug, Y-39983-HCl, which revealed dose-dependent phenotypic changes. At 1.0 μM, the drug caused significant alterations, including reduced cell size and irregular cell shape, aligning with its known disruption of actin filaments. The authors showed that a logistic regression model trained in a five-fold cross validation with plate held-out splits, was able to differentiate the drug from the controls with high accuracy (AUC = 1.0). At a lower dose of 0.2 μM, more subtle but still significant differences were observed, such as decreased nuclear eccentricity and altered cell granularity; this shows that lower doses can still induce phenotypic changes that are useful. We note that a leave-one-plate-out cross validation, where unique perturbations (compounds and dose) are spread across different plates, might leak information in most modelling approaches; however in this case, it is suitable for accounting for what makes a compound perturbation unique to the imaging across the experiment. Therefore, we retained the same leave-one-plate-out cross validation as the original authors.

We reproduced this experiment with only a single feature (“MitoCount”, the number of individual counts in the Mitochondria skeletonized mask, which is the closest related available feature to cell count) in order to compare their approach with a baseline cell count model. We found there was still a significant separation between the compound, Y-39983-HCl (1.0 μM), and controls when using the LR model with only MitoCount as a feature (AUC = 0.85), as shown in Figure 8b. For other compounds (including Y-39983-HCl at 0.2 μM), we observed better performance for ScaleFEx features (mean AUC = 0.99) compared to baseline models using MitoCount (mean AUC = 0.66). While the original paper identifies Zernike_ch1_1 as one of the features contributing to model performance and demonstrates its ability to distinguish Y-39983-HCl (1.0 μM) from the control, we additionally show that the MitCount feature achieves a comparable level of significance in this separation (Figure 8d). Overall, this shows that a baseline cell count-related model is a better benchmark than a random model. It also demonstrates that where phenotypic activity is subtle (at low concentrations), the full morphological profiles outperform a single feature related to cell count (as observed in the case of Y-39983-HCl at 0.2 μM dose).

**Figure 8.**
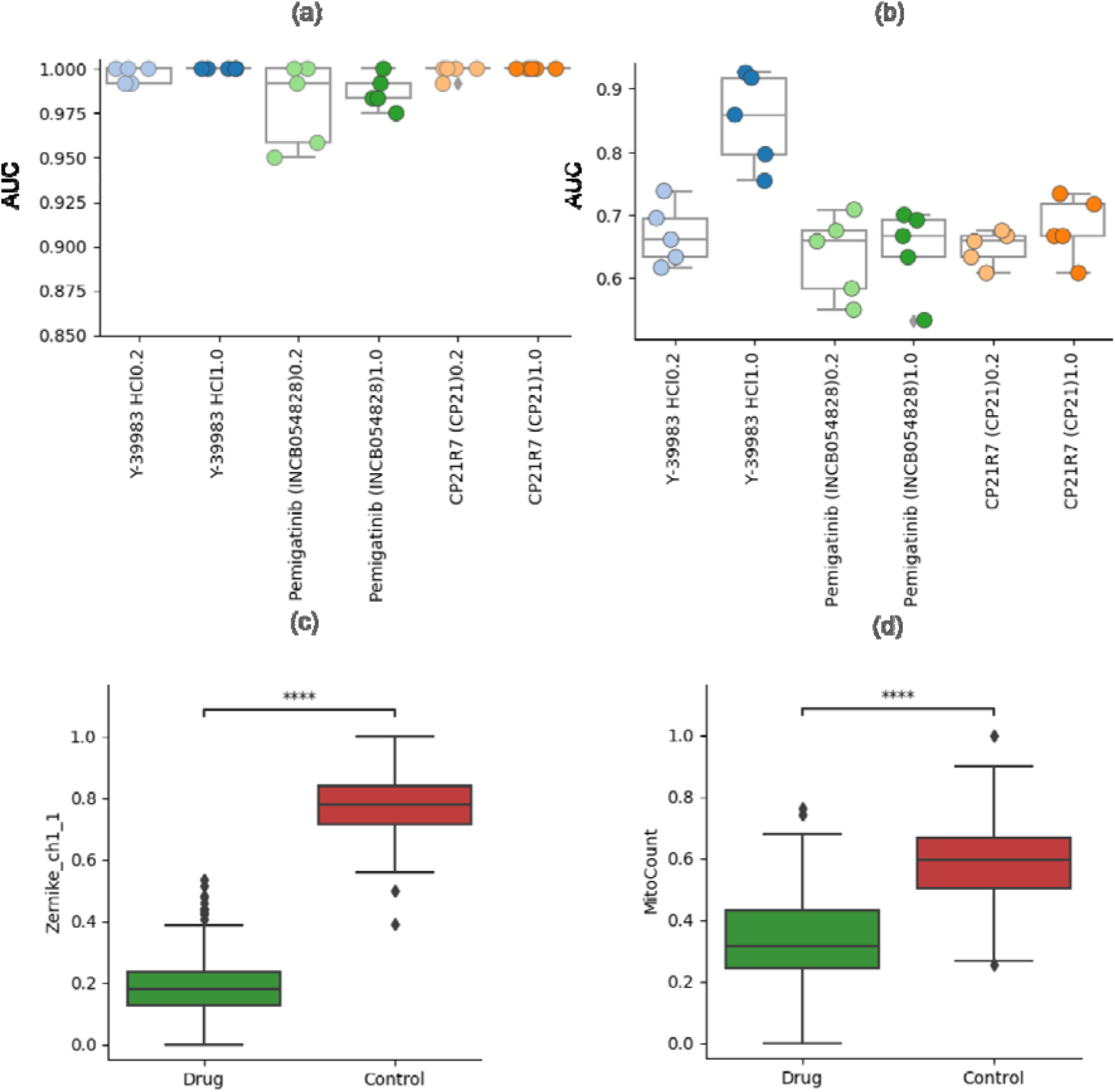
Distribution of AUC for (a) the model used in Comolet et al. and (b) the baseline cell count model employing the MitoCount feature, for a given drug versus the control, using a linear regression model with 5-fold cross-validation. The distribution of the most important feature value for correctly classifying Y-39983 HCl at 1.0 μM is shown for (c) the model from Comolet et al. and (d) the baseline cell count model with the MitoCount feature. Feature distribution values are normalized to a range of 0 to 1. Statistical significance is determined using a two-sided Mann–Whitney U test, with significance levels denoted as: ns (p > 0.05); ∗ (0.01 < p ≤ 0.05); and ∗∗∗∗ (p ≤ 0.0001).

To explore the concentration-dependency of information in Cell Painting profiles further, we evaluated dose-response dataset as a proof of concept (an earlier version of this dataset is described in Ewald et al^39^). We chose six compounds, chosen to highlight the separation between cytotoxic and bioactivity signals along concentration-response curves. Specifically, we chose three compounds that induced significant cell death at higher concentrations (Figure 9A-C). We also selected three compounds that exhibited distinct phenotypes without cytotoxicity across the tested concentration range (Figure 9D-F). Notably, the three cytotoxic compounds showed early morphological changes at concentrations ten times lower than those causing detectable cell death, as indicated by nucleus counts (Figure 9A-C). Although these findings are preliminary and specific to the six compounds, this is consistent with the analysis in Ewald et al., where they found that across >1000 compounds, morphology was perturbed about an order of magnitude lower than cell counts and other cytotoxicity readouts. This affirms that the information content in Cell Painting is dependent on dose.^39^

**Figure 9.**
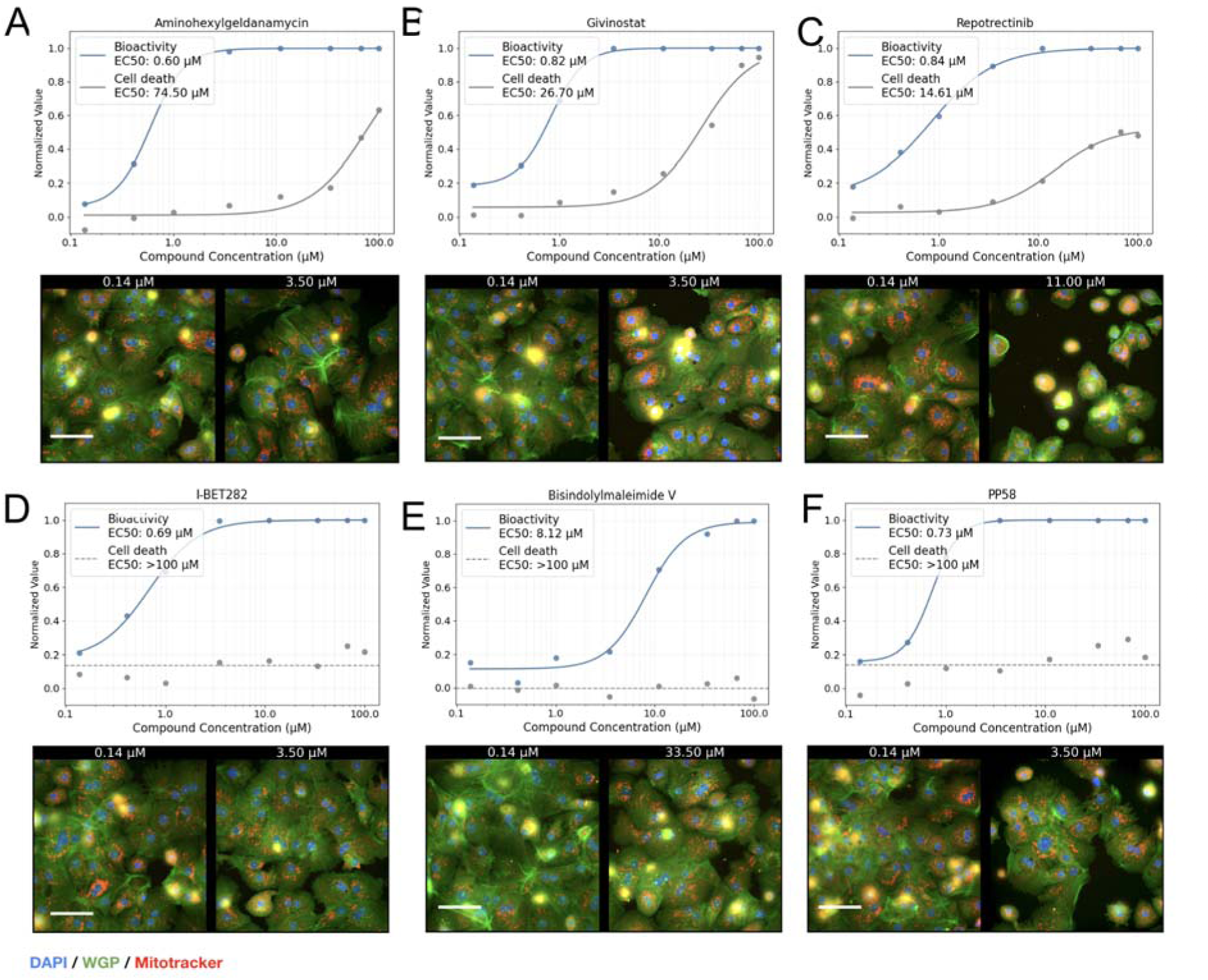
Image-based bioactivity screen with a viability screen for cytotoxicity. Bioactivity here refers to the morphological changes induced by compounds, measured as classifier probabilities distinguishing treated cells from DMSO controls. A probability of 0 indicates no detectable activity (indistinguishable from DMSO), while 1 represents significant morphological changes compared to DMSO. We show example dose-response curves for morphology change (bioactivity, blue), with higher values showing a perturbation being more dissimilar to DMSO, and cell death (nuclei count, gray), where higher values represent higher cell death as measured using nuclei count. A-C Example response curves for three cytotoxic compounds with various mechanisms of action (MOAs): A. Aminohexylgeldanamycin, an Hsp90 inhibitor B Givinostat, a histone deacetylase (HDAC) inhibitor, and C Repotrectinib, a tyrosine kinase inhibitor. Below each dose-response curve, representative 40x single field images for the lowest dose (0.14 µM) and first dose with very significant bioactivity (>0.9) are depicted. The scale bar is 80 µM. Cytotoxic compounds shrink the cytosol, and there are breaks in confluency. D-E Example response curves for compounds that were significantly bioactive but not did not considerably reduce nuclei count, including D. I-BET282 pan-inhibitor of all eight BET bromodomains, E. Bisindolylmaleimide V, protein kinase C inhibitor, and F PP58, Src Inhibitor.

### Recommendations for the Community

Following the above analyses, we make the following recommendations to best evaluate assay prediction models using Cell Painting and other high-dimensional phenotypic profiles such as gene expression:

1. **Prioritize assay selection to be aware of confounding influences in the benchmarking datasets.** The selection of assay endpoints is pivotal in ensuring reliable and interpretable results when benchmarking predictive models using Cell Painting or other high-dimensional phenotypic profiling techniques. Existing benchmarks are dominated by assays such as cytotoxicity or cell proliferation and risk obscuring the true predictive value of complex profiling features. This limitation is particularly pronounced in endpoints directly tied to cell viability, where the predictive utility of complex phenotypic profiles becomes marginal, and baseline models reliant on simple measurements, such as cell count, suffice. To address this, we recommend ensuring benchmarks contain assay endpoints that capture biologically nuanced processes or specific protein target interactions, where high-dimensional features offer genuine added value. Furthermore, curating diverse assay panels that do not over-represent viability-related tasks ensures balanced datasets, reduces bias in model evaluations, and enables more accurate assessments of complex phenotypic profiling methods.
2. **We recommend (a) analysing the distribution of cytotoxic compounds in each task, either using data from viability screens or cell count from Cell Painting assays (where available) and (b) looking at the images for a sample of correct (and confident) predictons.** Including cytotoxic compounds can skew results and produce misleading conclusions, predicting cell death instead of the intended biological signal. Many cytotoxicity-induced stress responses can activate cellular defense mechanisms, which may be detected in pathway-specific assays and erroneously interpreted as specific pathway activation by the test compound.^12^ Analysing the proportion of cytotoxic compounds could mitigate the confounding effects of including such biological assays in benchmarks. Further, it should be ensured that the selected assay is the correct summarization of the compound activities from the project and not the viability screen data, which may share the same assay project name in databases like PubChem. When using Cell Painting profiles, after models are developed and predictions made, we recommend viewing the raw images for evidence of cytotoxicity for a few of the correct and confident predictions. Determining whether all predictions are associated with images where most cells are dead can reveal if the model is learning to predict based on artifacts such as dark voids or low cell counts. The images from Bray et al. (https://idr.openmicroscopy.org/search) and the JUMP-CP project (https://phenaid.ardigen.com/jumpcpexplorer) are publicly searchable, providing resources to facilitate such examinations.^18,38^ For transcriptomic and proteomic profiling data types, this interpretation step would involve examining the particular genes contributing to successful predictions of an assay and evaluating whether they make biological sense or might be associated with an artifact or biological phenotype that is not of interest.
3. **Use baseline cell count models trained on a single cell count feature to compare with models trained on phenotypic profile data**. Baseline models that use cell count features offer a valuable benchmark for performance metrics, especially when a dataset contains assays directly linked to cell viability and/or when active compounds are disproportionately cytotoxic. We recommend training a baseline cell count model–a logistic regression model trained on the cell count feature (or a related feature like cell neighbour, mitocount etc.) but otherwise keeping the training/evaluation approach identical. Comparing with baseline models that predict outcomes solely based on cell count will reveal whether more complex models capture insights beyond cytotoxicity.
4. **Ensure a sufficient number of compounds with balanced labels and diversity in the test sets**. An insufficient number of test compounds, particularly with imbalanced classes, can lead to inflated performance metrics. For example, correctly predicting a single active compound in a test fold could misleadingly result in an AUC of 1. To mitigate this, Hofmarcher et al.^8^ ensured that each assay in their dataset included at least ten active and ten inactive measurements, maintaining this balance even after dataset splitting. For example, with models that achieve an AUC of 0.85, we recommend that held-out test sets contain at least 40 compounds from each class to be able to evaluate the model’s predictive performance robustly.^40^ When this is not feasible, AUC may not be an appropriate metric. Instead, we suggest evaluating models using absolute predictions instead of predicted probabilities, using metrics such as balanced accuracy, while carefully accounting for potential confounding effects from data imbalance using metrics such as AUPRG as well as random prediction baselines and y-scrambling.
5. **Recognize the biological context of the assay before model interpretation**. A general recommendation beyond cytotoxicity is that insight into the relevant biological mechanisms and pathways can elucidate the reasons behind a model’s performance. This involves recognizing typical cellular responses associated with the assay. We recommend interpreting the key features contributing to model performance ^41,42^; for instance, mitochondrial granularity features are more predictive of mitochondrial toxicity than features from other imaging channels.^19^ Additional factors influencing model interpretation include the choice of cell lines, treatment durations, and compound concentrations. Interpretation of image-based profiles, particularly those extracted by deep learning, is an area of active research.

### Benchmarking Protein-Target prediction with Cell Painting profiles compared cell counts

In order to facilitate above recommendations, we are together with this work releasing a benchmark dataset for compound-activity prediction curated along the recommendations above. This benchmark dataset contains bioactivity annotations for 1,349 compounds and 17 protein targets across 23 assays from DrugMatrix, PubChem and ChEMBL. This included a cytochrome panel, a secondary pharmacology panel, and binding affinity assays, comprising protein targets involved in signal transduction, neurotransmission, and metabolism, particularly with a focus on G protein-coupled receptor (GPCR) signaling, neurotransmitter activity, and xenobiotic metabolism (Supplementary Figure S1). We removed compounds that caused excessive cytotoxicity in the Cell Painting assay, and removed assays with fewer than 20 actives or 20 inactive compounds, resulting in a matrix with 50.8% completeness of which 27.24% were active compounds.

On this new benchmark, we found models using the complete Cell Painting profiles (mean balanced accuracy, BA=0.61, Figure 10a) are better at predicting assay outcome than the baseline cell count model (mean BA=0.51). When compared to the best of a random shuffling and a baseline cell count model, the models using complete Cell Painting profiles show an improvement of 30% or more in AUC-PR in 10 out of 23 tasks in the dataset (Supplementary Figure S2). Further, the improvements in AUC-ROC were consistent across the various sizes and the number/ratio of active compounds across all test sets, that is, confounding effects due to test set size are unlikely (Supplementary Figure S2).

**Figure 10.**
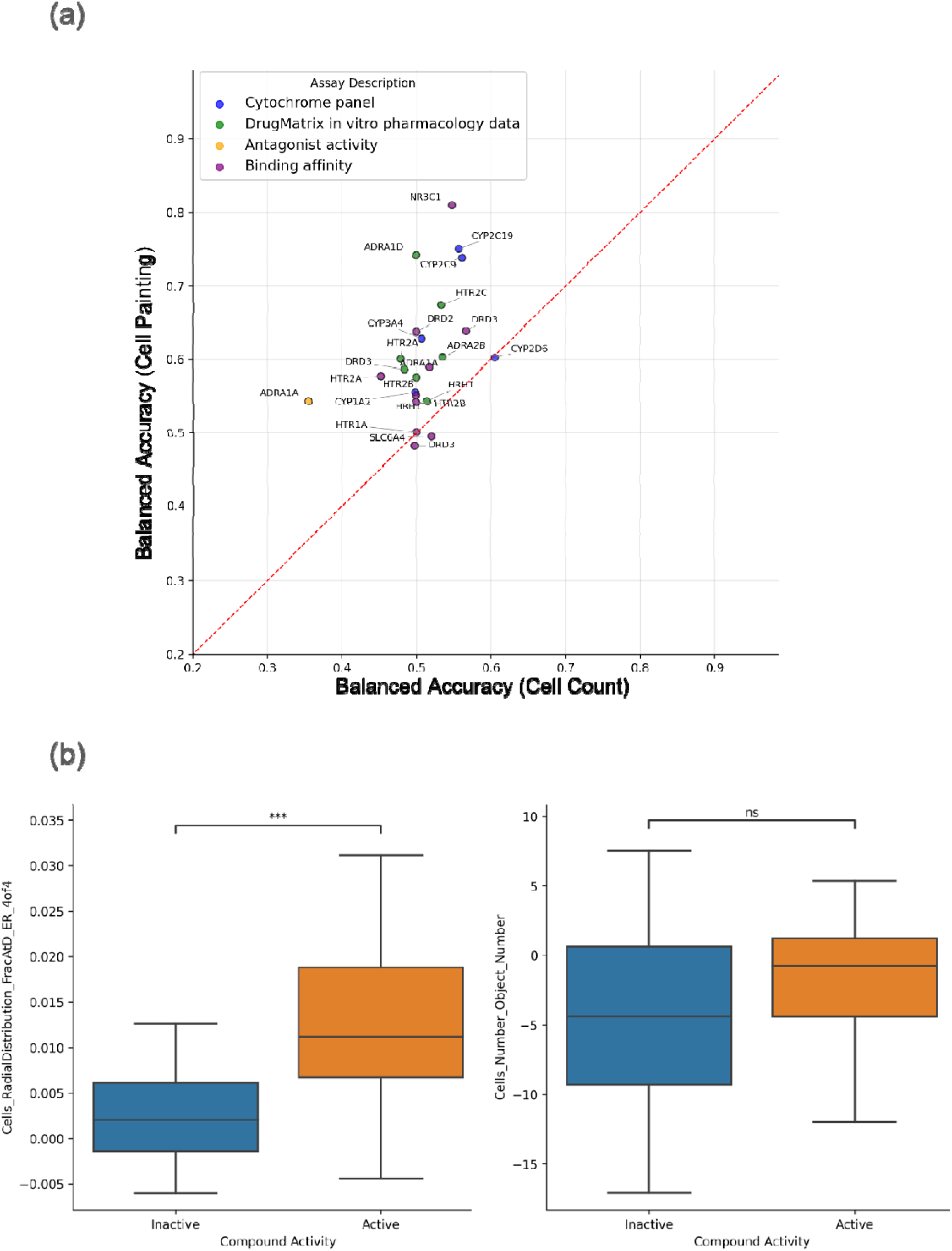
(A) Mean balanced accuracy for each of the 23 assays from evaluation of models using the complete Cell Painting profiles compared to the model using the cell count feature as a baseline. (B) Distribution the cell count and “Cells RadialDistribution FracAtD ER 4of4” features for active and inactive compounds in the NR3C1 binding assay. Statistical significance is determined using a two-sided Mann–Whitney U test, with significance levels denoted as: ns (p > 0.05); ∗∗∗ (0.0001<p ≤ 0.001).

The top performing assay, NR3C1 (glucocorticoid receptor) , is a proposed novel target for clear cell renal cell carcinoma.^43^ Nyffeler et al. have previously used Cell Painting to identify glucocorticoid receptor modulating chemicals.^44^ Given we found predictions from models trained for NR3C1 in this study are not driven by cell count (see Supplementary Figure S3), we hypothesized a model can separate the compounds binding to NR3C1 only with the full Cell Painting profiles. Our analysis indeed identified specific morphological and texture features as significant contributors to the prediction of NR3C1 binding activity. Features such as Cells_RadialDistribution_FracAtD_ER_4of4 (Figure 10b) and Cytoplasm_Texture_SumAverage_ER suggest alterations in the spatial distribution and organization of subcellular components, likely reflecting the receptor’s influence on the endoplasmic reticulum (ER) and protein folding machinery. Similarly, nuclear features such as Nuclei_AreaShape_Zernike and Nuclei_Texture_InfoMeas2 point to changes in nuclear morphology and chromatin texture, consistent with the receptor’s role as a transcription factor that translocates to the nucleus upon activation. The importance of mitochondrial and ER-related features aligns with NR3C1’s established role in regulating cellular stress responses and energy metabolism. Overall, these results underscore the utility of high-content imaging data for capturing biologically meaningful phenotypes associated with signaling pathways, not just limited to cell count.

## CONCLUSION

This study uncovers properties of existing assay prediction benchmarks that confound their goal of evaluating the predictive power of phenotypic profiles for diverse assay endpoints. We establish cell count as a baseline for high-dimensional profiling data, demonstrating that simple features are often highly predictive of assay outcomes. Particularly predictive for cell viability-related assays, these baselines serve as benchmarks to assess the added value of complete phenotypic profiles, as well as more complex model architectures, when predicting bioactivity. This limitation potentially biases the data towards compounds showing strong cytotoxic effects or drastic morphological changes, which are likely easier to detect and predict. We recommend five practices for using high-dimensional phenotyping readouts in machine learning tasks relevant to Cell Painting and other omics datasets–such as gene expression or protein expression datasets. Future studies could also explore if excluding cell-count-predictable assays during training (eg. multi-task neural networks) may improve model performance by preventing it from overfitting to cell count, though adversarial approaches could help remove confounding signals. We provide all code at https://github.com/srijitseal/The_Seal_Files (all data also released via https://doi.org/10.5281/zenodo.14838604)

## Acknowledgements

S. Seal acknowledges funding from the Cambridge Centre for Data-Driven Discovery (C2D3) Accelerate Programme for Scientific Discovery. S. Seal, S. Singh, and A.E.C. acknowledges funding from the National Institutes of Health (NIH MIRA R35 GM122547 to A.E.C.), the Massachusetts Life Sciences Center Bits to Bytes Capital Call program for funding the data production (to S. Singh), and the OASIS Consortium organised by HESI. O.S. acknowledges funding from the Swedish Research Council (grants 2020-03731, 2020-01865, 2024-03566, 2024-04576), FORMAS (grant 2022-00940), Swedish Cancer Foundation (22 2412 Pj 03 H), and Horizon Europe grant agreement #101057014 (PARC) and #101057442 (REMEDI4ALL). W. Dee acknowledges the UKRI/BBSRC Collaborative Training Partnership in AI for Drug Discovery, led by Exscientia Plc. in partnership with Queen Mary University of London. The Collaborative Training Partnership was funded by the Biotechnology and Biological Sciences Research Council, grant reference BB/X511791/1.

## Supplementary Information

**Figure S1.**
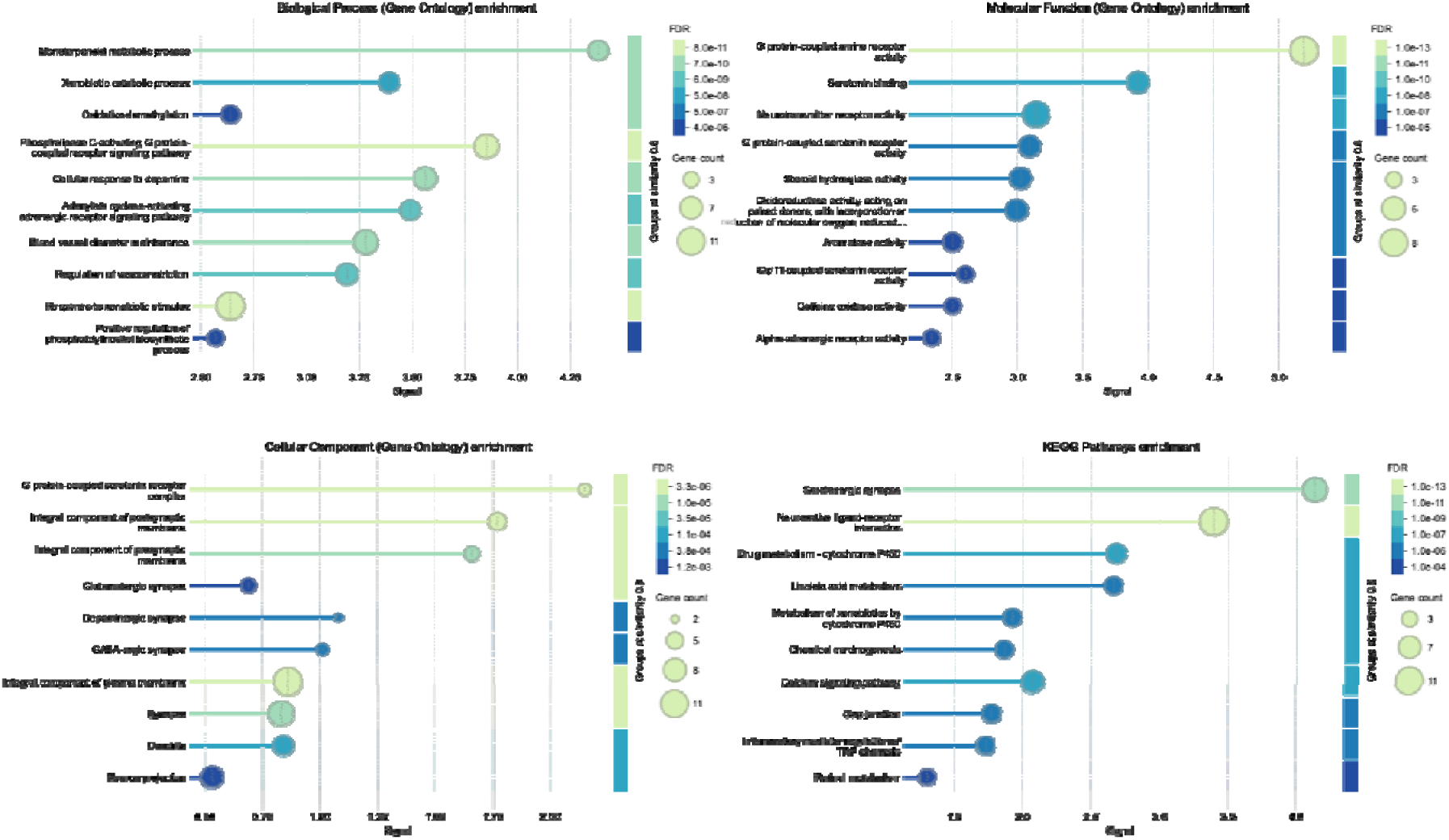
Characteristics of the dataset presented in this study, comprising 17 protein targets across 23 assays from DrugMatrix, PubChem and ChEMBL. Top 10 terms for functional enrichment visualization (similarity≥0.8) for (a) Biological process, (b) Molecular function, (c) Cellular component, and (d) KEGG pathways. Protein targets studies included a cytochome panel, secondary pharmacology targets and binding affinity, representing proteins involved in signal transduction, neurotransmission, and metabolism, particularly with a focus on G protein-coupled receptor (GPCR) signaling, neurotransmitter activity, and xenobiotic metabolism. (For details see https://version-12-0.string-db.org/cgi/network?networkId=b9rWbbS5Hlsu)

**Figure S2.**
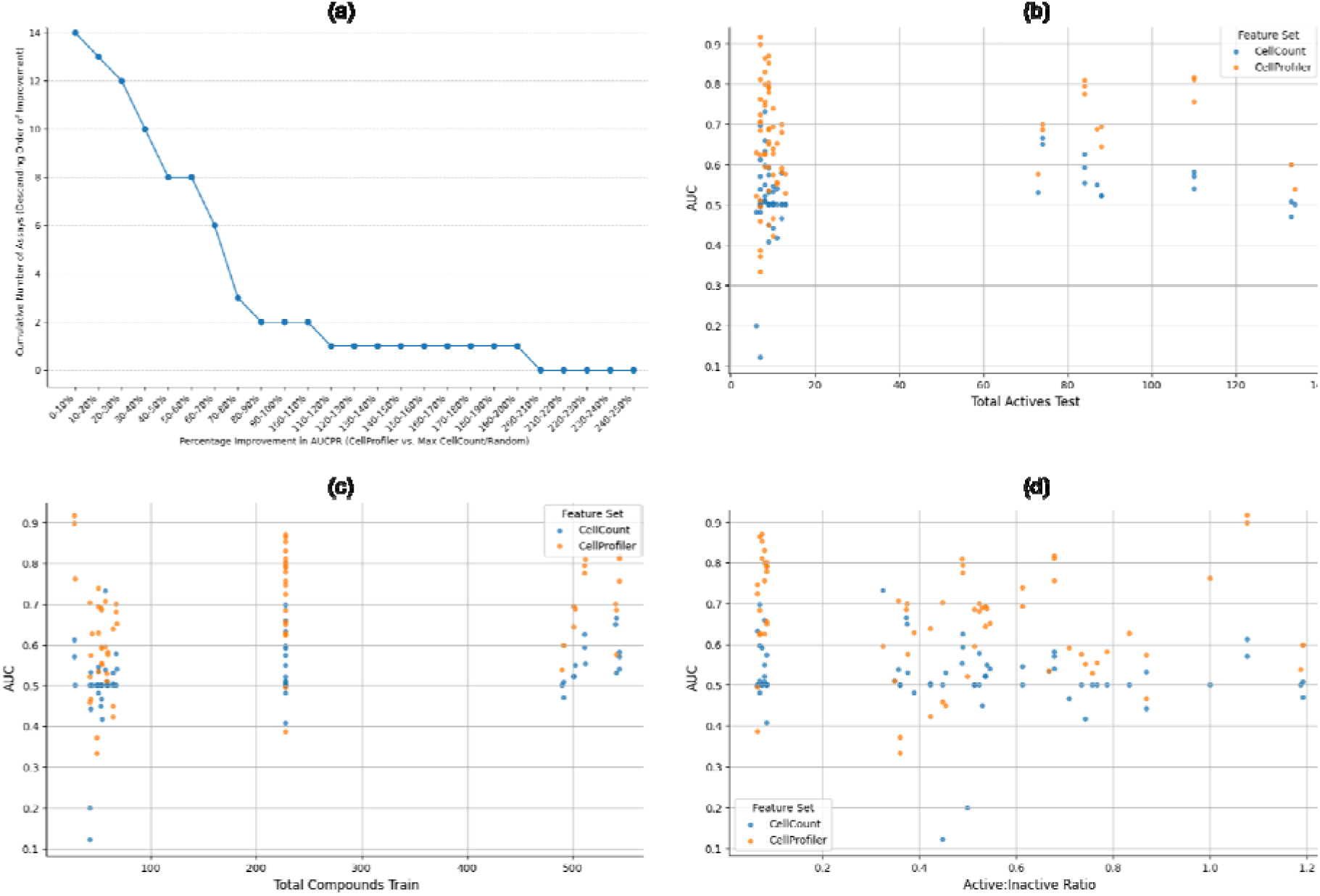
(a) Number of Assays out of 23 studied in this work where we observed a percentage improvement in AUCPR when using CellProfiler features compared to the best of a baseline cell count model of random shuffling or a baseline cell count model using only the cell count feature. Distribution of AUCROC achieved by model comapred to (b) total active compouines in the test dataset, (c) total compounds in the training data, and (d) the ratio of active to inactive compounds in the dataset. All dataset characteristics are mean of three folds, where given stratified splits result in individual characteristics of folds to be the same.

**Figure S3.**
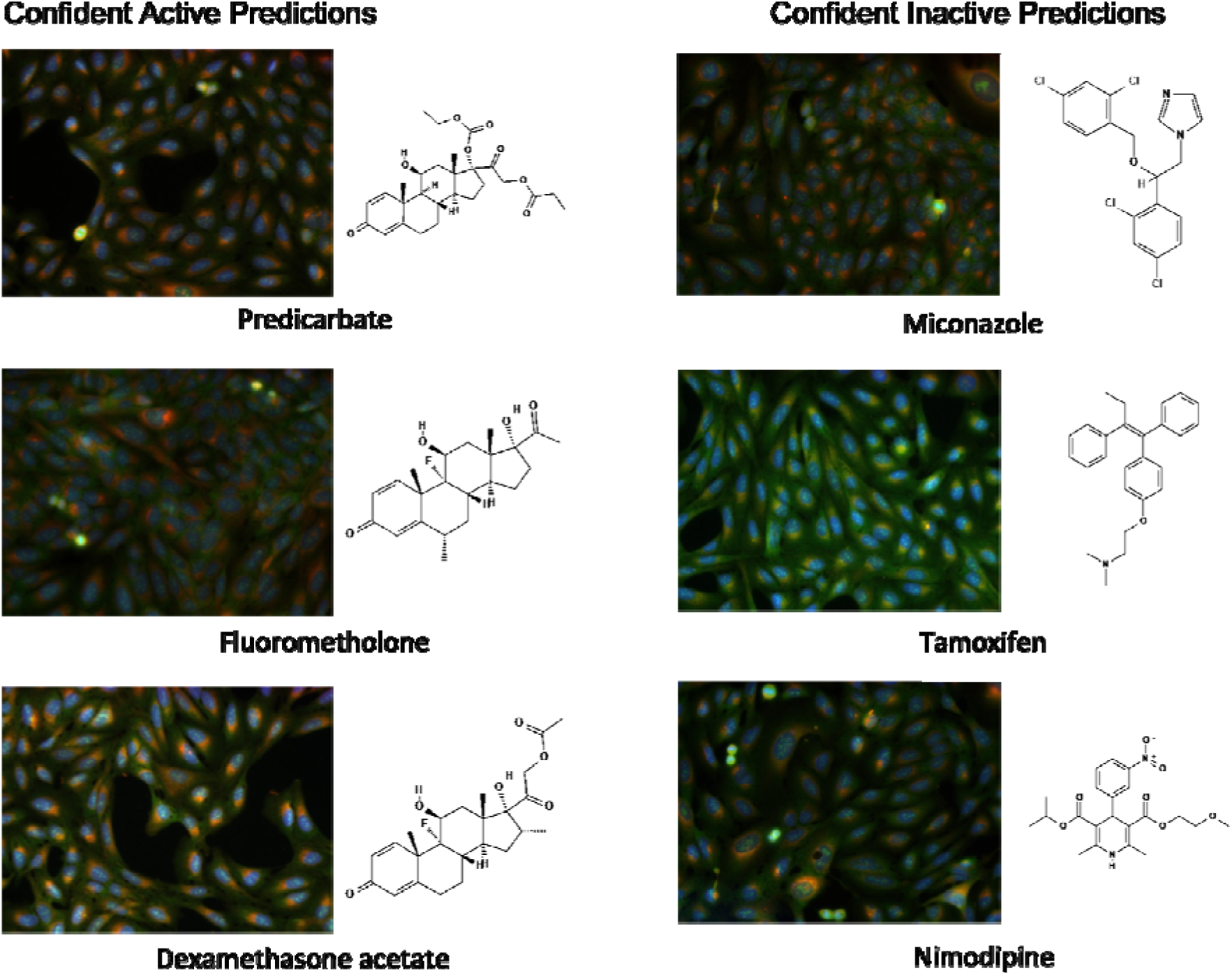
Predictions for NR3C1 binding affinity assay using complete Cell Painting profiles, correctly identified the glucocorticoid pharmacophore (fused four-ring steroid structure) from the imaging data. The positive predictions are known compounds that interact with the NR3C1 glucocorticoid receptor. No apparent changes in cell count is seen between active and inactive predictions, this indicating the Cell Painting profiles can learn more than cytotoxicity.

## REFERENCES

1. Bender, A. et al. Evaluation guidelines for machine learning tools in the chemical sciences. Nat Rev Chem 6, 428–442 (2022).

2. Zhou, H. & Skolnick, J. Utility of the Morgan Fingerprint in Structure-Based Virtual Ligand Screening. J. Phys. Chem. B 128, 5363–5370 (2024).

3. Hanser, T., Barber, C., Marchaland, J. F. & Werner, S. Applicability domain: towards a more formal definition. SAR QSAR Environ. Res. 27, 893–909 (2016).

4. Liu, A., Seal, S., Yang, H. & Bender, A. Using chemical and biological data to predict drug toxicity. SLAS Discov 28, 53–64 (2023).

5. Seal, S., et al. A Decade in a Systematic Review: The Evolution and Impact of Cell Painting. bioRxiv (2024) doi:10.1101/2024.05.04.592531.

6. Chandrasekaran, S. N., Ceulemans, H., Boyd, J. D. & Carpenter, A. E. Image-based profiling for drug discovery: due for a machine-learning upgrade? Nat. Rev. Drug Discov. 20, 145–159 (2021).

7. Simm, J. et al. Repurposing high-throughput image assays enables biological activity prediction for drug discovery. Cell Chem. Biol. 25, 611–618.e3 (2018).

8. Hofmarcher, M., Rumetshofer, E., Clevert, D.-A., Hochreiter, S. & Klambauer, G. Accurate Prediction of Biological Assays with High-Throughput Microscopy Images and Convolutional Networks. J. Chem. Inf. Model. 59, 1163–1171 (2019).

9. Moshkov, N. et al. Predicting compound activity from phenotypic profiles and chemical structures. Nat. Commun. 14, 1967 (2023).

10. Sanchez-Fernandez, A., Rumetshofer, E., Hochreiter, S. & Klambauer, G. CLOOME: contrastive learning unlocks bioimaging databases for queries with chemical structures. Nat. Commun. 14, 7339 (2023).

11. Ha, S. V., Leuschner, L. & Czodrowski, P. FSL-CP: a benchmark for small molecule activity few-shot prediction using cell microscopy images. Digital Discovery 3, 719–727 (2024).

12. Escher, B. I., Henneberger, L., König, M., Schlichting, R. & Fischer, F. C. Cytotoxicity Burst? Differentiating Specific from Nonspecific Effects in Tox21 in Vitro Reporter Gene Assays. Environ. Health Perspect. 128, 77007 (2020).

13. Dahlin, J. L. & Walters, M. A. The essential roles of chemistry in high-throughput screening triage. Future Med. Chem. 6, 1265–1290 (2014).

14. Ibrahim, M., Klindt, D. & Balestriero, R. Occam’s razor for Self Supervised Learning: What is sufficient to learn good representations? arXiv [cs.LG] (2024).

15. Bender, A. & Glen, R. C. A discussion of measures of enrichment in virtual screening: comparing the information content of descriptors with increasing levels of sophistication. J. Chem. Inf. Model. 45, 1369–1375 (2005).

16. Walters, P. Comparing Classification Models - You’re Probably Doing It Wrong. http://practicalcheminformatics.blogspot.com/2023/11/comparing-classification-models-youre.html (2023).

17. andreasbender. How to Lie With Computational Predictive Models in Drug Discovery. https://www.drugdiscovery.net/2020/10/13/how-to-lie-with-computational-predictive-models-in-drug-discovery/.

18. Bray, M.-A. et al. A dataset of images and morphological profiles of 30 000 small-molecule treatments using the Cell Painting assay. Gigascience 6, 1–5 (2017).

19. Seal, S. et al. Integrating cell morphology with gene expression and chemical structure to aid mitochondrial toxicity detection. Commun Biol 5, 858 (2022).

20. Statistical functions (scipy.stats) — SciPy v1.11.3 Manual. https://docs.scipy.org/doc/scipy/reference/stats.html.

21. Flach, P. A. & Kull, M. Precision-Recall-Gain curves: PR analysis done right. Neural Inf Process Syst 838–846 (2015).

22. Cross-Zamirski, J. O. et al. Label-free prediction of cell painting from brightfield images. Sci. Rep. 12, 10001 (2022).

23. Comolet, G., et al. A highly-efficient, scalable pipeline for fixed feature extraction from large-scale high-content imaging screens. bioRxiv (2023) doi:10.1101/2023.07.06.547985.

24. Rücker, C., Rücker, G. & Meringer, M. y-Randomization and its variants in QSPR/QSAR. J. Chem. Inf. Model. 47, 2345–2357 (2007).

25. Pedregosa, F. et al. Scikit-learn: Machine Learning in Python. arXiv [cs.LG] 2825–2830 (2012).

26. PubChem. AID 651635 - viability Counterscreen for Primary qHTS for Inhibitors of ATXN expression - PubChem. https://pubchem.ncbi.nlm.nih.gov/bioassay/651635.

27. PubChem. AID 624202 - qHTS Assay to Identify Small Molecule Activators of BRCA1 Expression - PubChem. https://pubchem.ncbi.nlm.nih.gov/bioassay/624202.

28. Saito, T. & Rehmsmeier, M. The precision-recall plot is more informative than the ROC plot when evaluating binary classifiers on imbalanced datasets. PLoS One 10, e0118432 (2015).

29. Varalda, M. et al. Psychotropic Drugs Show Anticancer Activity by Disrupting Mitochondrial and Lysosomal Function. Front. Oncol. 10, 562196 (2020).

30. Wang, C.-C., Chen, B.-K., Chen, P.-H. & Chen, L.-C. Hinokitiol induces cell death and inhibits epidermal growth factor-induced cell migration and signaling pathways in human cervical adenocarcinoma. Taiwan. J. Obstet. Gynecol. 59, 698–705 (2020).

31. Kim, J.-H., Chae, M., Choi, A.-R., Sik Kim, H. & Yoon, S. SP600125 overcomes antimitotic drug-resistance in cancer cells by increasing apoptosis with independence of P-gp inhibition. Eur. J. Pharmacol. 723, 141–147 (2014).

32. Bray, M.-A. et al. Cell Painting, a high-content image-based assay for morphological profiling using multiplexed fluorescent dyes. Nat. Protoc. 11, 1757–1774 (2016).

33. Luo, Y. et al. Characterization and analysis of biopharmaceutical proteins. in Handbook of Modern Pharmaceutical Analysis 283–359 (Elsevier, 2011).

34. Schmitt, D. M. et al. The use of resazurin as a novel antimicrobial agent against Francisella tularensis. Front. Cell. Infect. Microbiol. 3, 93 (2013).

35. Liu, J. et al. Wnt/β-catenin signalling: function, biological mechanisms, and therapeutic opportunities. Signal Transduct. Target. Ther. 7, 3 (2022).

36. Chen, C. et al. Aberrant activation of Wnt/β-catenin signaling drives proliferation of bone sarcoma cells. Oncotarget 6, 17570–17583 (2015).

37. Zhang, Y. & Wang, X. Targeting the Wnt/β-catenin signaling pathway in cancer. J. Hematol. Oncol. 13, 165 (2020).

38. Chandrasekaran, S. N. et al. JUMP Cell Painting dataset: morphological impact of 136,000 chemical and genetic perturbations. bioRxiv 2023.03.23.534023 (2023) doi:10.1101/2023.03.23.534023.

39. Ewald, J. D. et al. Cell Painting for cytotoxicity and mode-of-action analysis in primary human hepatocytes. bioRxiv 2025.01.22.634152 (2025).

40. Hanley, J. A. & McNeil, B. J. The meaning and use of the area under a receiver operating characteristic (ROC) curve. Radiology 143, 29–36 (1982).

41. Seal, S. et al. From pixels to phenotypes: Integrating image-based profiling with cell health data as BioMorph features improves interpretability. Mol. Biol. Cell 35, mr2 (2024).

42. Pahl, A. et al. Morphological subprofile analysis for bioactivity annotation of small molecules. Cell Chem Biol 30, 839–853.e7 (2023).

43. Yan, M. et al. Knockdown of NR3C1 inhibits the proliferation and migration of clear cell renal cell carcinoma through activating endoplasmic reticulum stress-mitophagy. J. Transl. Med. 21, 701 (2023).

44. Nyffeler, J. et al. Application of Cell Painting for chemical hazard evaluation in support of screening-level chemical assessments. Toxicol. Appl. Pharmacol. 468, 116513 (2023).

